# Differential physiological role of BIN1 isoforms in skeletal muscle development, function and regeneration

**DOI:** 10.1101/477950

**Authors:** Ivana Prokic, Belinda Cowling, Candice Kutchukian, Christine Kretz, Hichem Tasfaout, Josiane Hergueux, Olivia Wendling, Arnaud Ferry, Anne Toussaint, Christos Gavriilidis, Vasugi Nattarayan, Catherine Koch, Jeanne Lainné, Roy Combe, Laurent Tiret, Vincent Jacquemond, Fanny Pilot-Storck, Jocelyn Laporte

## Abstract

Skeletal muscle development and regeneration are tightly regulated processes. How the intracellular organization of muscle fibers is achieved during these steps is unclear. Here we focus on the cellular and physiological roles of amphiphysin 2 (BIN1), a membrane remodeling protein mutated in both congenital and adult centronuclear myopathies, that is ubiquitously expressed and has skeletal muscle-specific isoforms. We created and characterized constitutive, muscle-specific and inducible *Bin1* homozygous and heterozygous knockout mice targeting either ubiquitous or muscle-specific isoforms. Constitutive *Bin1*-deficient mice died at birth from lack of feeding due to a skeletal muscle defect. T-tubules and other organelles were misplaced and altered, supporting a general early role of BIN1 on intracellular organization in addition to membrane remodeling. Whereas restricted deletion of *Bin1* in unchallenged adult muscles had no impact, the forced switch from the muscle-specific isoforms to the ubiquitous isoforms through deletion of the in-frame muscle–specific exon delayed muscle regeneration. Thus, BIN1 ubiquitous function is necessary for muscle development and function while its muscle-specific isoforms fine-tune muscle regeneration in adulthood, supporting that BIN1 centronuclear myopathy with congenital onset are due to developmental defects while later onset may be due to regeneration defects.

## Introduction

Skeletal muscle is composed of bundles of multinucleated myofibers containing dozens to thousands of nuclei in a common cytoplasm. Specific organelles as the myofibril contractile apparatus and the triads sustain contraction and force development (Franzini-Armstrong 2018). The triad sustains the excitation-contraction (EC) coupling machinery and is composed of a plasma membrane invagination (T-tubule) connecting two terminal cisternae from the sarcoplasmic reticulum (SR). The membrane network is exquisitely developed in muscle with the SR wrapping the myofibrils (Towler et al. 2004). During muscle development in mammals, other organelles achieve specific localization: for e.g. mitochondria concentrate around myofibrils while nuclei are positioned at the periphery of the fiber. Underneath the basal lamina that surrounds each myofiber are satellite cells (Mauro 1961) which are the muscle stem cells implicated in muscle growth and regeneration. In mammals, several steps of muscle remodeling occur: muscle development before birth, maturation around and shortly after birth, muscle maintenance, and muscle regeneration upon injury (Engel and Franzini-Armstrong 2004). Muscle regeneration recapitulates to a certain extent the developmental process of muscle formation. Satellite cells proliferate in response to growth factors produced by muscle injury and fuse with existing myofibers. A plethora of muscle diseases impair different steps of muscle formation or maintenance (Demonbreun and McNally 2016). Hence, understanding the key players in these processes represents a crucial step towards the development of therapeutic approaches. Here we investigated the physiological role of BIN1, a protein mutated in rare myopathies, on muscle development, function and regeneration.

BIN1, or amphiphysin 2, is a main regulator of membrane remodeling and trafficking (McMahon and Gallop 2005; Itoh and De Camilli 2006; Prokic et al. 2014), and its role in organelle positioning begins to emerge (Falcone et al. 2014; D’Alessandro et al. 2015). It belongs to the amphiphysin protein family which N-terminal BAR domain is known to recognize and induce membrane curvature (Peter et al. 2004; Frost et al. 2009). It also has a SH3 domain binding to proline-rich motifs in other proteins. BIN1 is a ubiquitously expressed protein highly expressed in skeletal muscles and brain (Sakamuro et al. 1996; Butler et al. 1997). The *BIN1* gene has 20 exons and encodes multiple isoforms. Skeletal muscle expresses several isoforms including ubiquitous isoforms that contain both the BAR and C-terminal SH3 domains and muscle-specific isoforms that in addition contain the inframe exon 11 (named exon 10 in previous studies on the cDNA) encoding a polybasic motif binding phosphoinositides (PIs)(Lee et al. 2002; Fugier et al. 2011). The PI+ muscle-specific isoforms appear during muscle development and are the prevalent isoforms in adult ((Cowling et al. 2017) and this study). Overexpression of the muscle-specific BIN1 iso8 (a PI+ isoform) but not ubiquitous (PI-) isoforms in C2C12 myoblasts and other cells results in the formation of numerous tubules connected to the plasma membrane, suggesting the PI domain promotes membrane remodeling (Lee et al. 2002; Nicot et al. 2007; Bohm et al. 2013). In addition to binding phosphoinositides, the muscle-specific PI motif regulates a conformational switch by binding the SH3 domain of BIN1 molecules (Kojima et al. 2004; Royer et al. 2013). The abundance of PIs such as PtdIns(4,5)P_2_ may regulate this conformational switch and therefore the binding to downstream interactors of the SH3 domain, such as dynamin or synaptojanin (Kojima et al. 2004).

*BIN1* is mutated in different forms of centronuclear myopathies (CNM), characterized by muscle hypotrophy, muscle weakness, centralized nuclei and triad defects (Romero 2010; Toussaint et al. 2011; Jungbluth and Gautel 2014; Prokic et al. 2014). Considering recessive CNM cases with nonprogressive muscle involvement, homozygous missense mutations in the BAR domain impair the membrane tubulation properties while premature truncation of the SH3 domain reduces the binding to dynamin 2 (Nicot et al. 2007). Moreover, a splice site mutation leading to the skipping of in-frame exon 11 was identified in humans and dogs presenting with highly progressive CNM (Bohm et al. 2013). Exon 11 skipping was also associated to myotonic dystrophies (DM), characterized by muscle wasting and increased fiber degeneration and regeneration (Fugier et al. 2011). In addition, heterozygous BIN1 mutations cause an adult-onset mild CNM (Bohm et al. 2014). In all cases, the primary affected organ is the skeletal muscle but how *BIN1* mutations in different domains lead to different onset and severity is not understood. Findings in cells and animal models suggested a role of BIN1 on the formation of T-tubules (Razzaq et al. 2001; Lee et al. 2002; Tjondrokoesoemo et al. 2011; Bohm et al. 2013; Smith et al. 2014). However, the physiological role of BIN1 in mammalian muscle remains elusive.

In this study, we characterized several novel *Bin1* knockout (KO) mice. In *Bin1* KO mice deleted for exon 20 (*Bin1*x20-/-), the SH3 domain found in all isoforms was disrupted, similarly as in some CNM patients with truncating mutations, which makes these mice a good model to study the ubiquitous function of BIN1. In *Bin1* KO mice missing exon 11 (*Bin1*x11-/-), the muscle-specific isoforms were converted into ubiquitous isoforms by deletion of this in-frame exon. This later model helped address the muscle-specific function of PI+ BIN1 isoforms. Molecular, cellular and physiological characterization of constitutive, muscle-specific and inducible lines uncovered that while BIN1 is necessary for skeletal muscle development and function at birth, its muscle-specific isoforms are dispensable for development but required for muscle regeneration in adult.

## Results

### *Bin1* has the highest expression in skeletal muscle, where it is mainly located on the triads

In adult humans, *BIN1* is ubiquitously expressed with the highest expression in skeletal muscle (Figure S1A)(Sakamuro et al. 1996; Butler et al. 1997; Nicot et al. 2007; GTEx_consortium 2015). To address *Bin1* expression during mouse embryonic development we performed in situ hybridization at E14.5 and E18.5 using a probe against the 3’UTR region of *Bin1* (Figure 1A). The highest expression of *Bin1* was found in skeletal muscle and diaphragm, followed by the cortex and the eye. To assess the localization of muscle-specific BIN1 isoforms in skeletal muscle, immunogold labeling was performed in the adult *tibialis anterior* (TA) muscle using an anti-BIN1 antibody directed against the PI domain encoded by the muscle-specific exon 11. This staining revealed that muscle-specific BIN1 specifically localizes at the triads but not to other membrane structures or compartments (Figure 1B). These findings suggested an important role of BIN1 in muscle development for the formation of triads.

**Figure 1.**
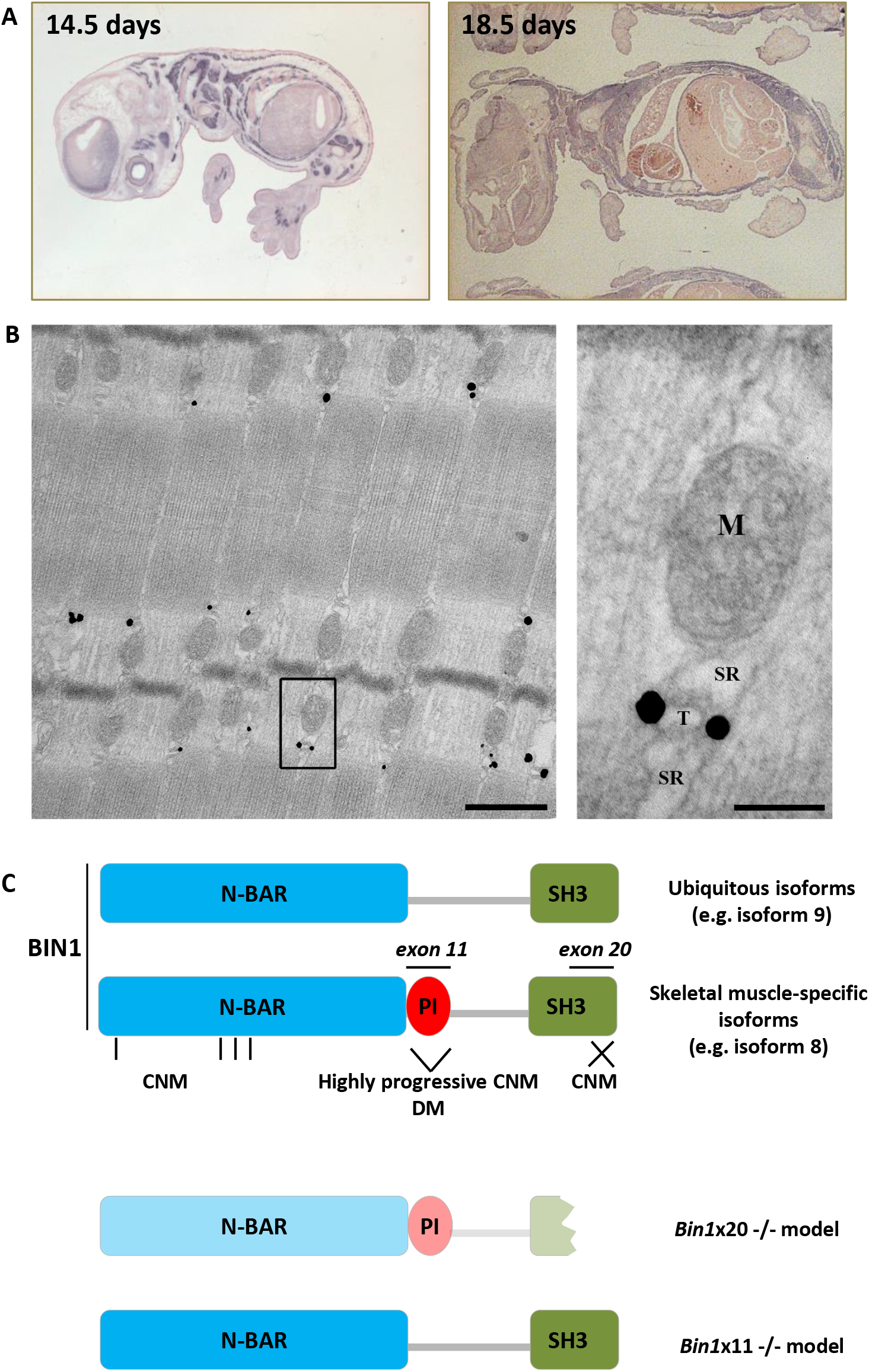
BIN1 expression and localization. (A) *In situ* BIN1 labeling in WT mouse at embryonic day 14.5 (E14.5) and 18.5 (E18.5). (B) Immunogold labeling using anti-PI domain specific antibody on a *tibialis anterior* muscle from a WT adult mouse showing specific triad localization. SR=sarcoplasmic reticulum, T=T-tubule, M=mitochondria. Scale bars are 500 nm and 100 nm respectively. (C) BIN1 functional domains encompass the N-BAR (N-terminal amphipathic helix with Bin-Amph-Rvs), the PI (phosphoinositides binding), and the SH3 (src homology) domains. Exon 11 encodes the muscle specific PI domain while exon 20 encodes the second half of the ubiquitous SH3 domain. Missense mutations in the N-BAR or truncation of the SH3 domains lead to recessive centronuclear myopathy (CNM) while skipping of the PI domain is linked to highly progressive CNM and myotonic dystrophy (DM). *Bin1*x20-/- mice express a strongly reduced and truncated BIN1 while *Bin1*x11-/- mice express the ubiquitous isoforms in muscle.

### Constitutive and muscle-specific homozygous deletion of *Bin1* exon 20 leads to feeding inability and perinatal death

To address the physiological importance of BIN1 and generate a BIN1-CNM model, we created and analyzed *Bin1*x20-/- mice, deleted in the last exon 20 (Figure 1C) and thus causing disruption of the SH3 domain similarly to some CNM patients (Figure S1B-C)(Cowling et al. 2017). The complete homozygous deletion of *Bin1* is lethal in the first hours after birth and cardiomyopathy was proposed to be the fatal cause (Muller et al. 2003). However, mice with a cardiac-specific deletion of *Bin1* survived (Laury-Kleintop et al. 2015). To further clarify the death-causing mechanism of *Bin1* deficiency, we analyzed survival. We confirmed that this constitutive deletion induced precocious lethalithy with no *Bin1*x20-/- mice surviving the first postnatal day among 131 littermates. Homozygous deletion of *Bin1* exon 20 led to a strong decrease in the protein level, and there was no compensatory overexpression of its close homolog amphiphysin 1 (Figure S1D-E). Comparison of *Bin1*x20-/- and WT littermates showed neither difference in body weight nor in heart organization and functioning (Figure S2A-E).

We then used the Cre recombinase under the control of the human skeletal actin promoter to induce the deletion specifically in skeletal muscle. As in the case of constitutive KO, the muscle-specific homozygous deletion led to a strong BIN1 reduction with no *Bin1*x20skm-/- mice surviving the first hours after birth (Figure 2; Figure S1F). A muscle-based lethality could originate either from breathing or feeding defect. HE staining of the diaphragm from *Bin1*x20-/- mice showed that organization and thickness did not differ from control mice (Figure 2A). Accordingly, their lungs inflated and no cyanosis was observed, suggesting a functional diaphragm. Conversely, HE analysis of the *quadriceps* revealed increased centralization of nuclei, a hallmark of CNM (Figure 2A-B). Even several hours after birth and unlike their WT littermates, constitutive *Bin1*x20-/- mice had an empty stomach (Figure 2C). They also had a very low glucose level (Figure 2D), supporting strong hypoglycemia as the cause of death. To exclude a prenatal hypoglycemia, blood glucose levels were measured at E18.5 and were similar to WT (Figure 2E). Similar findings were observed in the *Bin1*x20skm-/- mice (Figure 2F-G). Altogether, these results show that BIN1 is necessary for muscle development and function, and for proper feeding after birth.

**Figure 2.**
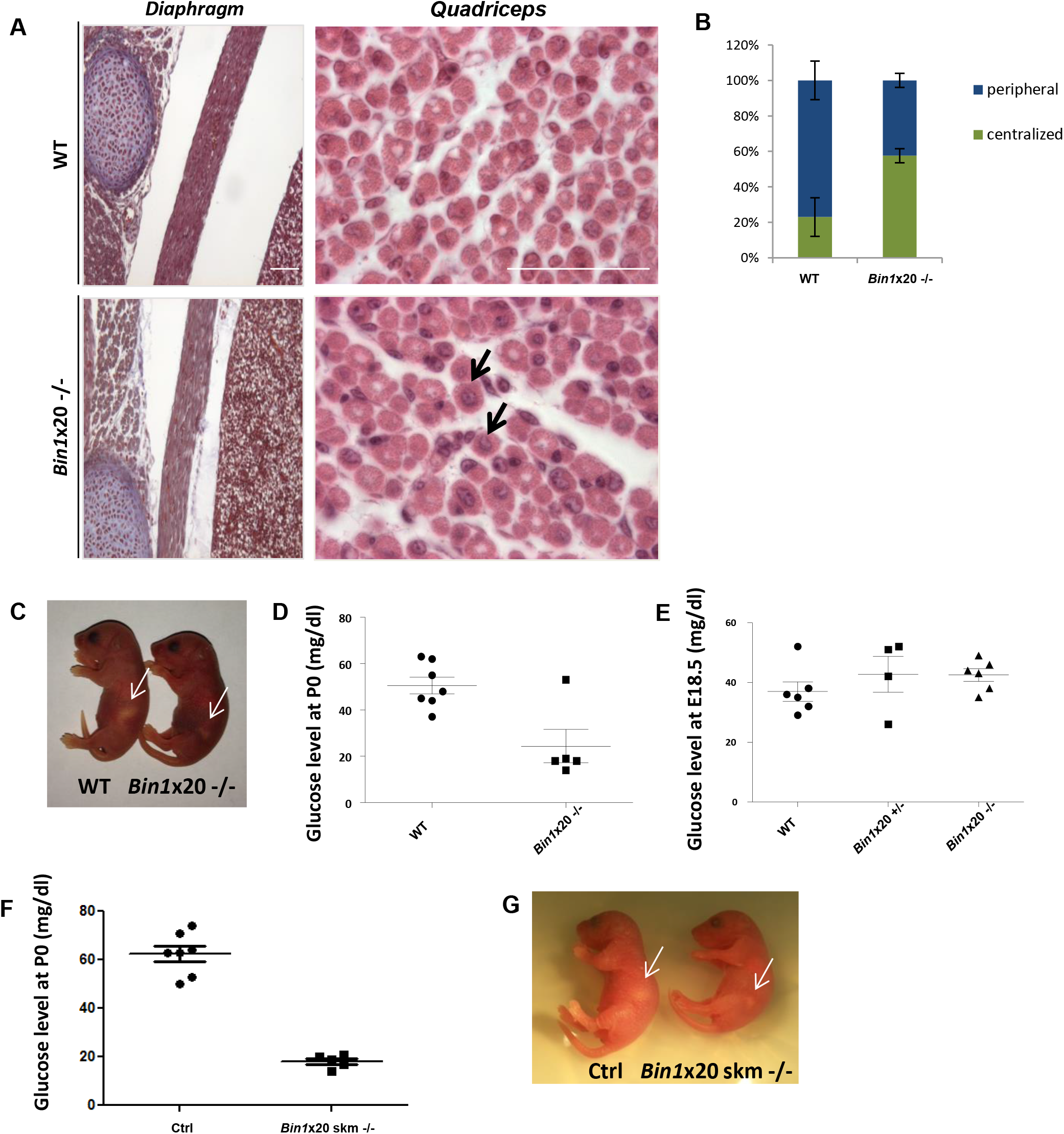
Muscle-related lethality in *Bin1*x20-/- mice. (A) Histology of the diaphragm and quadriceps in newborn WT and *Bin1*x20-/- mice. Scale bar 200 μm. (B) Quantification of centralized nuclei (arrows) versus peripheral in quadriceps. (C) Blood glucose level in E18.5 embryos showing no difference in the glucose level between the WT, *Bin1*x20+/- and constitutive *Bin1*x20-/- mice (n ≥ 4 mice per group). (D) Blood glucose level in P0 newborns. Most *Bin1*x20-/- mice showed a 3 fold reduction in glucose level compared to WT littermates (n ≥ 5 mice per group). (E) *Bin1*x20-/- mice had an empty stomach (arrows) compared to WT littermates. (F) Glucose measurement at P0 of muscle specific *Bin1*x20skm-/- showed a 3-fold reduction compared to WT littermates. (G) *Bin1*x20skm-/- mice have an empty stomach compared to WT littermates.

### BIN1 is necessary for triad formation and general muscle fiber organization

To better understand the cellular defects due to BIN1 alteration, we performed histology, ultrastructure and immunolabeling analyses. NADH-TR and SDH staining of *Bin1*x20-/- muscle sections showed strong collapse of the oxidative activity towards the center of fibers (Figure S3A-B). Alteration in mitochondria distribution was confirmed using an antibody against prohibitin, a mitochondria protein. Ultrastructure by TEM confirmed the presence of amorphous material and collapsed mitochondria and nuclei in the center of fibers that was deprived of myofibrils (Figure 3A), reminiscent of a CNM-like histology (Toussaint et al. 2011).

**Figure 3.**
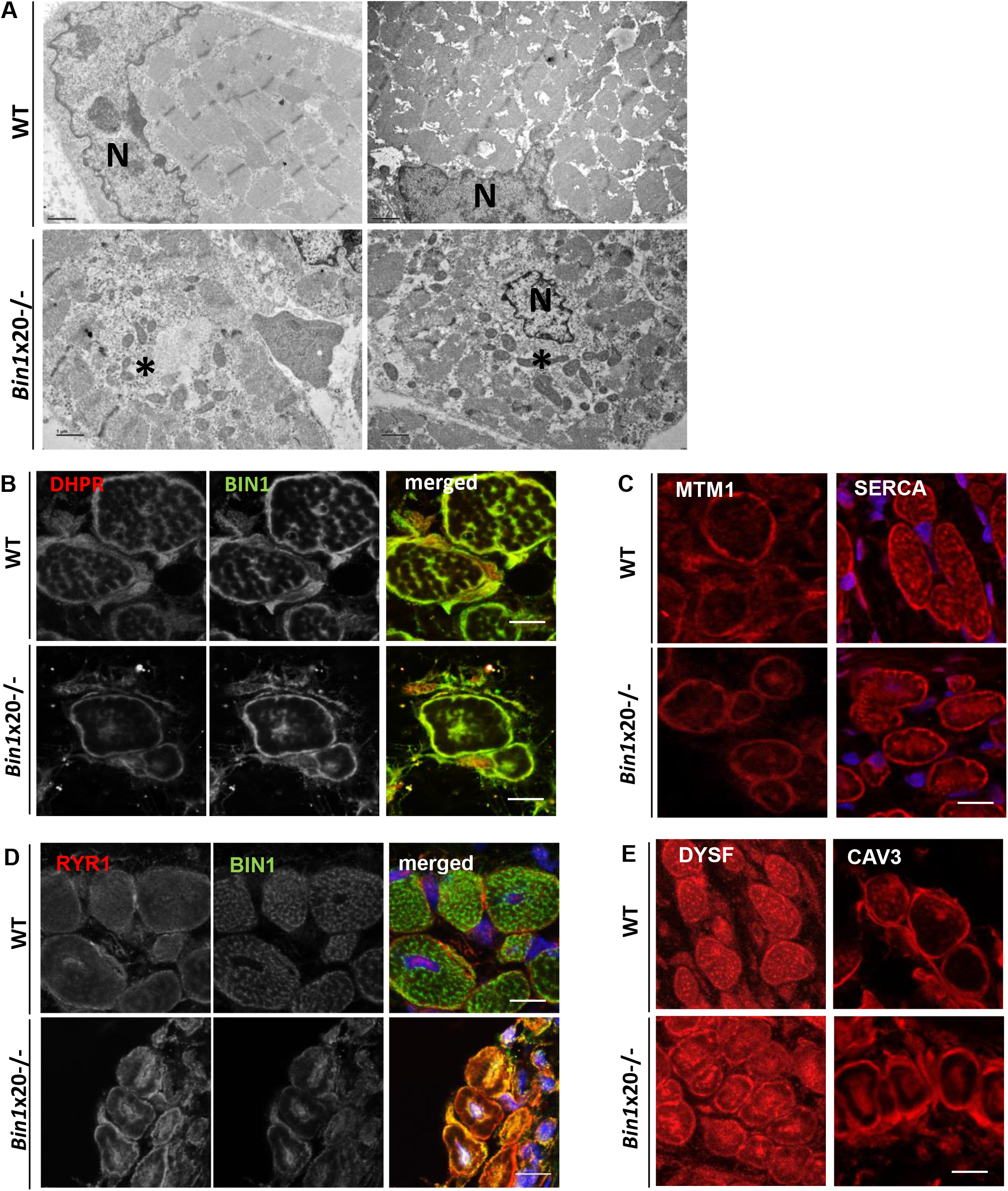
Alterations in intracellular organization and triads upon BIN1 defect. (A) EM ultrastructure showed a general disorganization of muscle from newborn *Bin1*x20-/- mice with a central collapse of nuclei (N) and surrounded by an area devoid of myofibrils and filled with mitochondria and amorphous materials (*). Scale bar 1 μm. (B) BIN1 detected with a pan-isoform antibody colocalized with the T-tubule marker DHPR in adult muscle fiber (transversal view). Markers of T-tubules (DHPR, B), junctional sarcoplasmic reticulum (MTM1 in C, RYR1 in D), longitudinal sarcoplasmic reticulum (SERCA in C) were collapsed to the center of myofibers in the *Bin1*x20-/- mice. (E) Markers of premature T-tubules (dysferlin and caveolin 3) were collapsed to the center in *Bin1*x20-/- mice. Note that muscle fibers in WT newborn are usually less polygonal than in adult. Immunolabeling of 8 μm transversal section. Scale bar 10 μm.

In muscle transversal sections, BIN1 colocalized with the T-tubule channel DHPR in WT while residual truncated BIN1 was barely visible and aggregated in *Bin1*x20-/-, along with DHPR (Figure 3B). In addition, the SR markers RYR1, SERCA1 and MTM1 were also mislocalized in *Bin1*x20-/-muscles (Figure 3C-D), supporting a general alteration of triads. Unlike triad markers, localization of sarcomeric markers α-actinin and DNM2 was not changed (Figure S4A-B).

To investigate if triad alterations was primarily due to a defect in T-tubule formation, we first localized in muscle sections dysferlin and caveolin 3, two proteins implicated in T-tubule formation, and found them also mislocalized, similarly to DHPR (Figure 3E). Second, we validated that BIN1 localized as longitudinal tubules in isolated fibers, a pattern lost in *Bin1*x20-/- (Figure 4A). DHPR and MTM1 were also abnormally localized in these isolated fibers (Figure 4B-C). Third, to assess the organization of T-tubules in formation, the FM4-64 impermeable lipophilic dye was applied to differentiated primary myotubes from WT and *Bin1*x20-/- mice. While the T-tubules network was dense and developed in WT myotubes, it was strongly reduced in *Bin1*x20-/- (Figure 4D). These data support a primary defect in T-tubule formation, leading to alteration of the triad in mature muscle and impairment of organelles and myofibrils positioning.

**Figure 4.**
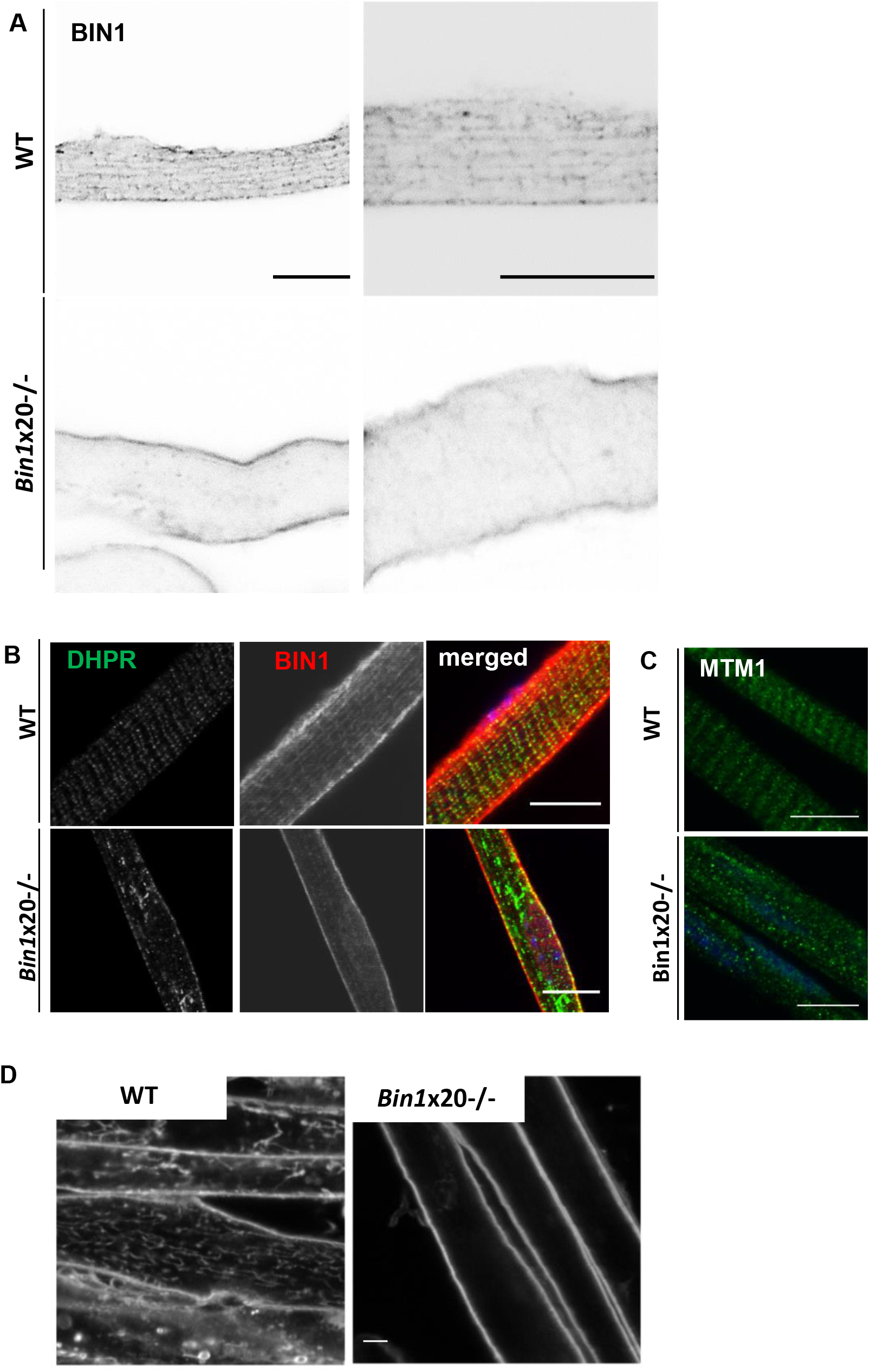
Defects in T-tubule and triad formation. (A) BIN1 detected with a pan-isoform antibody localizes on intracellular longitudinal tubules in newborn isolated myofibers, a pattern lost in *Bin1*x20-/- mice. (B) DHPR aggregation in *Bin1*x20-/- isolated myofibers. (C) MTM1 localization was disrupted in *Bin1*x20-/- isolated myofibers. (D) Primary differentiated myotubes with FM4-64 staining of sarcolemma-connected membrane tubules; *Bin1*x20-/- myotubes lack the dense longitudinal tubules network. Scale bar 5 μm.

### BIN1 is dispensable for adult muscle maintenance

To assess if the effects of *Bin1* exon 20 deletion were dose-dependent, *Bin1*x20+/- heterozygous mice were scrutinized. Heterozygous mice were found at the expected Mendelian ratio, had a normal lifespan and showed no obvious differences compared to their WT littermates. Body and muscle weight were comparable to WT (Figure S5A-B). *Bin1* heterozygous deletion did not affect the muscle performance assessed by the grip and rotarod tests (Figure S5C-D). Muscle histology was normal as well as the T-tubules organization (Figure S5E-F). Thus, BIN1 defect does not have a dose-dependent effect in skeletal muscle, unlike in heart where it causes T-tubule defects and increasing susceptibility to ventricular arrhythmias (Hong et al. 2014).

Since deletion of *Bin1* exon 20 impacted on perinatal muscle development, we also investigated BIN1 importance in adult muscle maintenance. For this, *Bin1* exon 20 floxed mice were crossed with mice expressing a tamoxifen-inducible Cre recombinase under the control of the actin promoter to restrict *Bin1* deficiency specifically in adult muscle. Following tamoxifen treatment at 7 weeks, *Bin1*x20skm(i)-/- mice were analyzed after 5 and 25 weeks. *Bin1* mRNA expression was strongly decreased after 5 w with a 20% decrease in BIN1 protein level that reached 80% reduction at 25 w (Figure 5; Figure S6). Exon 20 deletion was skeletal muscle specific and did not impair BIN1 cardiac expression (Figure 5A). We conducted thorough investigation at 25 w and found no difference in body weight, muscle mass, grip, string and hanging tests in *Bin1*x20skm(i)-/- mice (Figure S7). Histology with HE and NADH-TR, and electron microscopy revealed no differences in fiber crosssectional area or muscle organization (Figure 5B-D). Taken together, these data showed that BIN1 is dispensable for adult muscle maintenance under unchallenged conditions.

**Figure 5.**
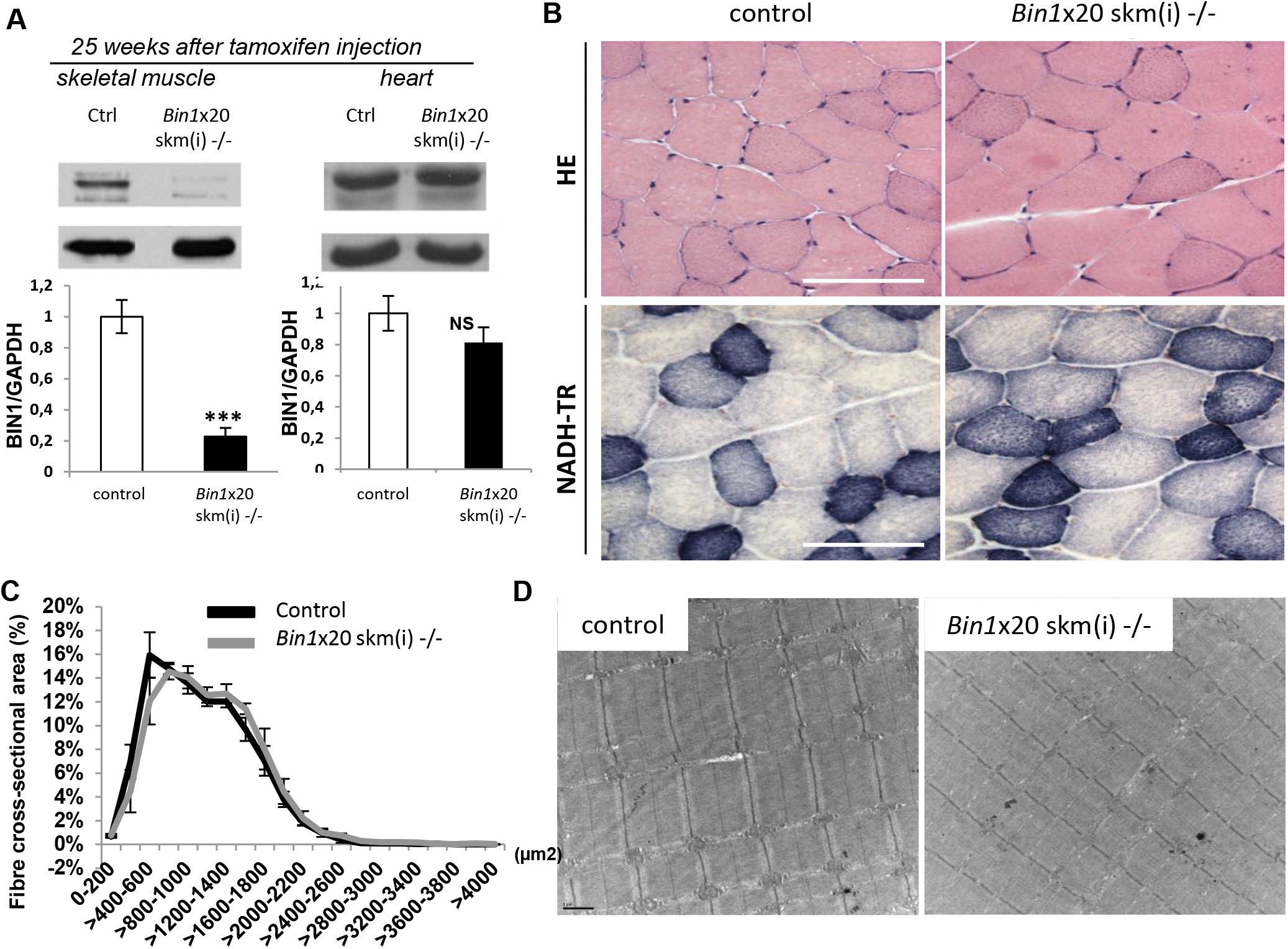
BIN1 is not essential for adult muscle maintenance. Characterization of *Bin1*x20skm(i)-/- mice 25 weeks after tamoxifen injection. (A) BIN1 protein level 25 weeks after intraperitoneal tamoxifen injection. Graph depict mean ± s.e.m. (***p < 0.001). Student t-test; n ≥ 3 mice per group. (B) Histology of TA muscles stained with hematoxylin and eosin (HE, upper panel) and NADH-tetrazolium reductase (NADH-TR, lower panel). Scale bar 100 μm. (C) Fiber areas of TA muscles were grouped into 200 μm^2^ intervals, and represented as the percentage of total fibers in each group (n ≥ 5 mice). (D) Electron microscopy on TA muscles. Scale bar 1 μm.

### Removal of BIN1 muscle-specific isoforms does not affect muscle development and function

To understand the particular function of BIN1 muscle-specific isoforms encompassing exon 11 in muscle development and maintenance, we constitutively deleted this in-frame exon to obtain *Bin1*x11-/- mice expressing only ubiquitous BIN1 isoforms in skeletal muscle and mimicking *BIN1* exon 11 skipping found in myotonic dystrophy and in the highly progressive form of CNM (Fugier et al. 2011; Bohm et al. 2013). Specific excision of exon 11 in muscle, without alteration in the splicing of neighboring exons, was confirmed by RT-PCR and sequencing of the corresponding cDNAs (Figure S8A-D). Note that adult muscle contains mainly BIN1 PI+ isoforms (iso8 +/- exon 17). Western blot from muscle extracts using an antibody against the PI domain (encoded by exon 11) detected no signal (Figure S8E). Other BIN1 isoforms were still detected with a pan-antibody raised against the SH3 domain. Removal of *Bin1* muscle-specific isoform caused no detectable upregulation of amphiphysin 1 (Figure S8F).

*Bin1*x11-/- mice were born following the expected Mendelian ratio and had normal lifespan, supporting that the PI domain is dispensable for muscle development. Phenotyping young adults at 12 weeks showed no difference with WT in body weight and length, muscle weight or fat and lean mass distribution (Figure S9A-D). Deletion of exon 11 neither impacted motor performance, as measured using grip and rotarod tests, nor reduced specific muscle force (Figure S10A-D). Muscle histology (HE and SDH stainings), fiber cross-sectional area and muscle ultrastructure were comparable between *Bin1*x11-/- and WT mice (Figure 6A-C; Figure S11). Only nuclei mis-localization was slightly but significantly increased from 1% in WT to 3.7% in *Bin1*x11-/- animals in the TA (Figure S11B). We also looked if loss of muscle-specific BIN1 isoforms had any impact on T-tubule and triad organization. BIN1 without the PI domain still localized to the triad. Immunofluorescence and potassium ferrocyanide revealed normal T-tubules structure and organization in *Bin1*x11-/- muscle (Figure 6B-C).

**Figure 6.**
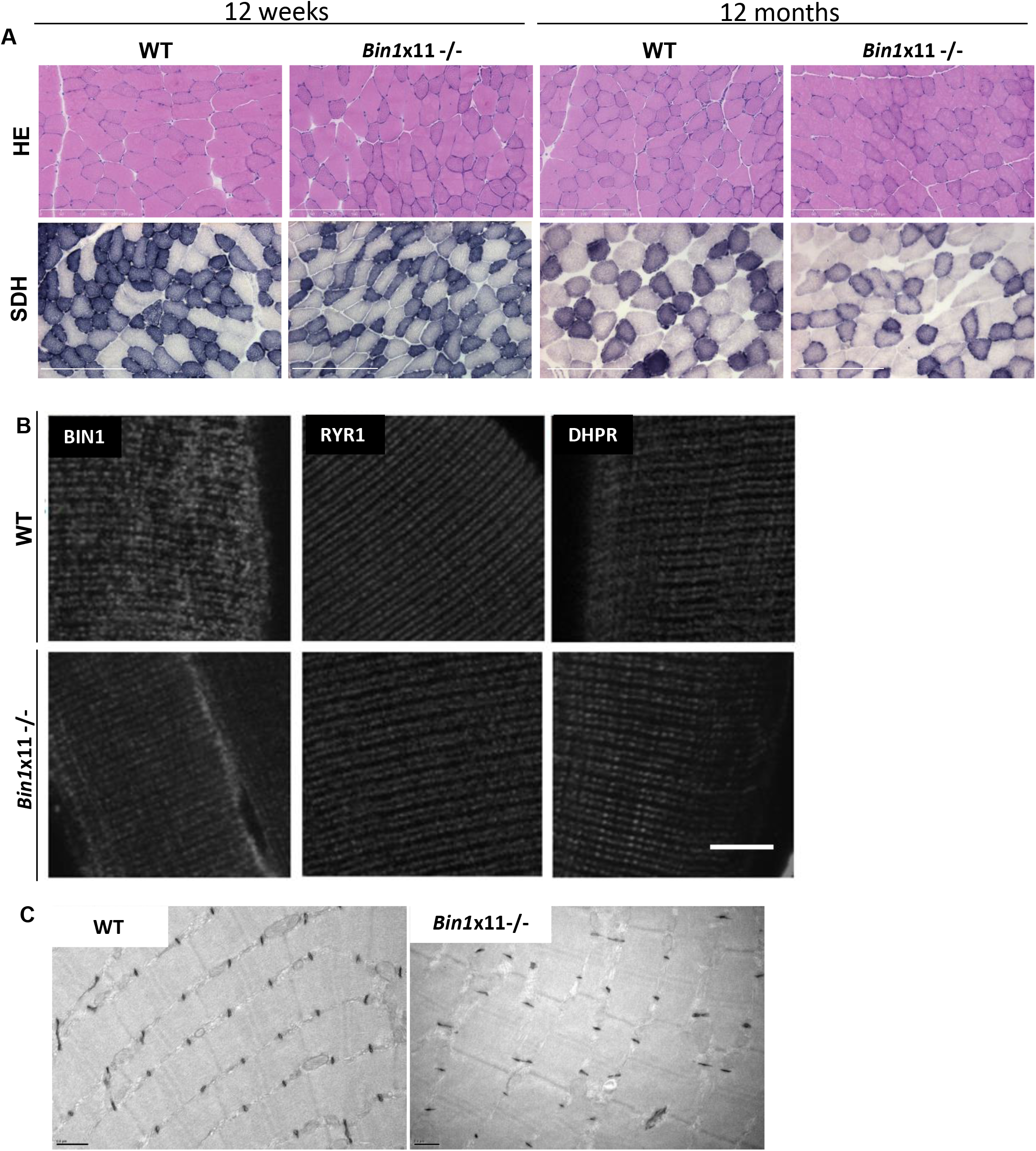
Lack of BIN1 muscle-specific isoforms correlates with normal histology and triad structure. (A) Histological features in *Bin1*x11-/- mice at 12 weeks and 12 months of age with hematoxylin and eosin (HE) or succinate dehydrogenase (SDH). Scale bar 200 μm. (B) Localization of BIN1 without the PI domain and triad markers in isolated fibers stained with BIN1 (pan-isoform antibody), RYR1 (sarcoplasmic reticulum), or DHPR (T-tubules) antibodies. Scale bar 10 μm. (C) Electron micrograph of 12-week-old WT and *Bin1*x11-/- muscles labeled with potassium ferrocyanide for T-tubules. Scale bar 0.5 μm.

Di-8-anepps staining revealed a slight reduction of the T-tubule network density (Figure 7), although in average mean values for membrane capacitance of the portion of fiber under study did not differ between fibers from *Bin1*x11-/- and WT mice (1.45 ± 0.09 nF, n=19 fibers, *versus* 1.59 ± 0.13 nF, n = 18 fibers, respectively). To assess in more details whether there was a functional defect of the excitation-contraction coupling in *Bin1*x11-/- mice, we analyzed functional properties of the DHPR and of the ryanodine receptor (RYR1) in intact isolated muscle fibers. Analysis of the voltage-dependence of DHPR Ca^2+^ current density revealed no change in maximum conductance, apparent reversal voltage, steepness factor and half-activation voltage in the *Bin1*x11-/- group (Figure S12). Regarding RYR1 activity, analysis of the voltage-dependence of the peak rate of SR calcium release revealed no significant difference in maximum amplitude, time to peak, voltage for half-activation and steepness factor between WT and *Bin1*x11-/- fibers (Figure S13). Voltage-activated Ca^2+^ transients were also measured with the low affinity dye fluo-4 FF in the absence of exogenous intracellular Ca^2+^ buffer, to assess the myoplasmic Ca^2+^ removal capabilities of the fibers. Fitting a single exponential function to the decay of the fluo-4 FF transients showed no difference between mean values in the two groups, suggesting no change in Ca^2+^ buffering and SR calcium uptake rate by SERCA pumps (Figure S14). Overall, the PI domain appears dispensable for myofibers structure and function.

**Figure 7.**
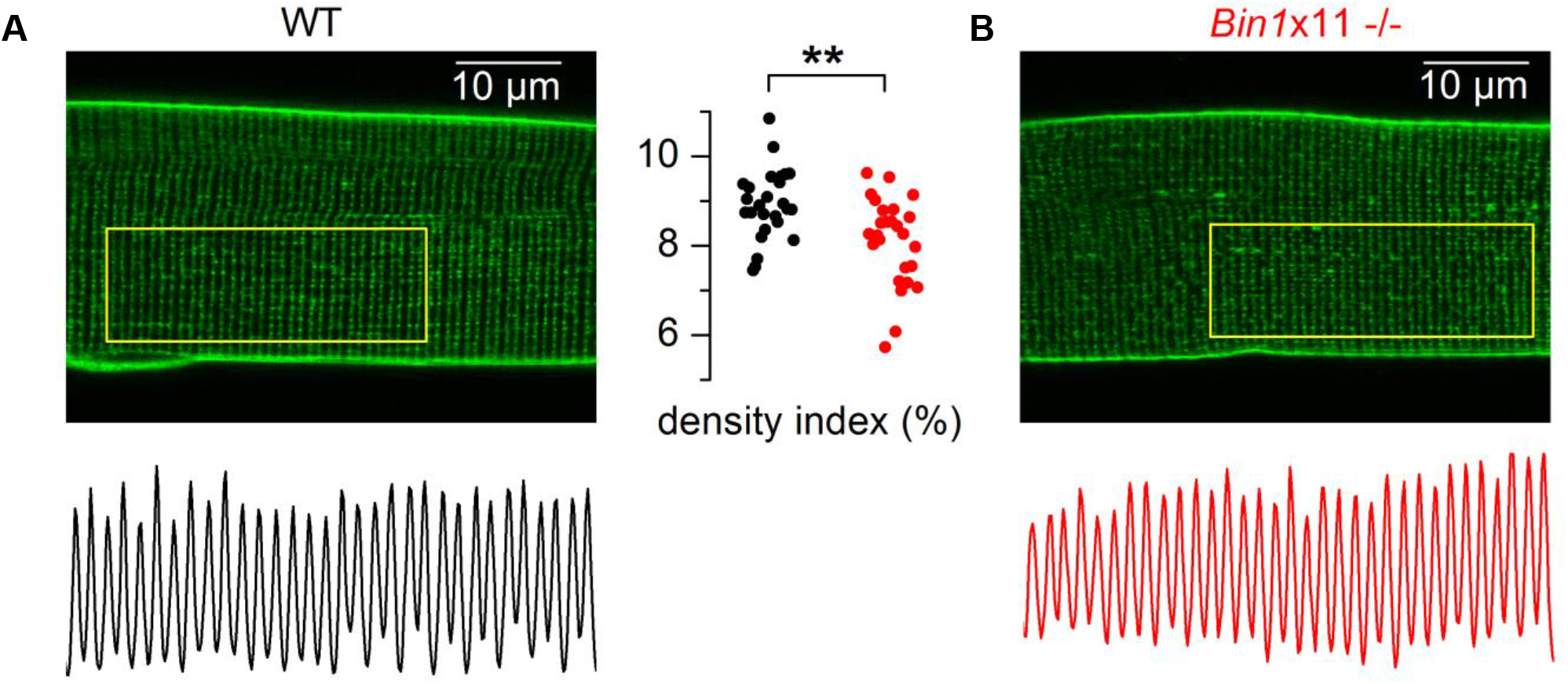
T-tubule network in muscle fibers from WT and *Bin1*x11-/- mice. *x,y* fluorescence images of di-8-anepps staining in a fiber from a WT mouse (A) and from a *Bin1*x11-/- mouse (B). The longitudinal profile of fluorescence along the outlined region of interest is shown below each image. The graph in the middle shows values for the T-tubule density index in fibers from WT mice (black points) and *Bin1*x11-/- mice (red points).

Longterm deletion (12 months of age) of exon 11 caused no additional motor, histological, ultrastructural anomalies or triad defects in *Bin1*x11-/- mice (Figure 6A; Figure S11B, S15 and S16). Mouse muscles mature until 7 weeks of age (Engel and Franzini-Armstrong 2004). To see if there were any early defects during postnatal muscle maturation in *Bin1*x11-/- mice, we looked at 2 weeks after birth. Body weight and muscle mass were comparable to WT (Figure S17A-B). Histology revealed no anomaly, and there was no difference in fiber size, nuclei localization, sarcomere and T-tubule organization (Figure S17C-F).

In conclusion, while BIN1 ubiquitous isoforms are mainly implicated in muscle development, the muscle-specific isoforms appear dispensable for muscle formation and maintenance raising the question of their specific function.

### The BIN1 muscle-specific isoforms are beneficial for muscle regeneration

The only phenotype observed in *Bin1*x11-/- mice in young and old mice was a slight increase in nuclei centralization, a feature observed during normal muscle regeneration. To test the ability of *Bin1*x11-/- muscle to regenerate, TA muscles were injected with notexin to produce muscle damage and analyzed 3, 5, 7, 14, 21 and 28 days after the injury. Normalization of the recovery of the damaged muscle towards the contralateral control muscle showed that, at the later stages of regeneration (14 and 28 days), the weight of the damaged *Bin1*x11-/- muscle significantly failed to recover compared to WT (Figure 8A). Histological examinations and quantification confirmed it was correlated to smaller fiber cross-sectional area (Figure 8B-C; Figure S18). The specific muscle force showed a lag in the recovery at 28 days, supporting that muscles without BIN1 muscle-specific isoforms produced intrinsically less force than WT muscles at late stage of muscle regeneration (Figure 8D).

**Figure 8.**
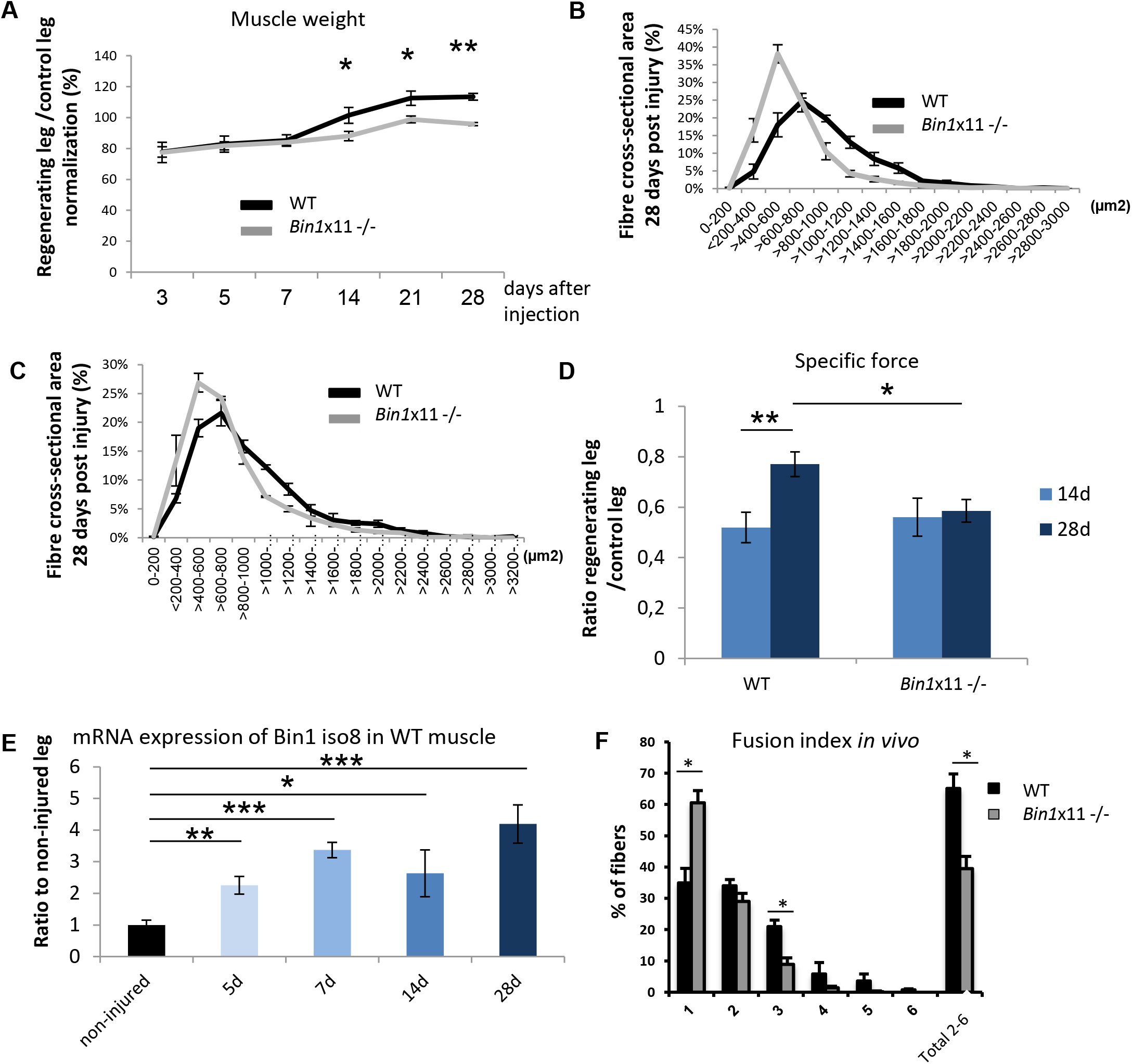
Muscle regeneration is altered in *Bin1*x11-/- mice. Right TA muscle of 12 weeks mice were injected with notexin and left TA used as control. (A) Regenerative capacity, estimated through normalization of the muscle mass of injected leg to uninjected contralateral leg at different time points; represented as the percentage of recovery. TA muscles fiber area at 14 days (B) and 28 days (C) after notexin injection. Only regenerating fibers (with centralized nuclei) were taken into account and fiber cross-sectional area is grouped into 200 μm^2^ intervals and represented as the percentage of total fibers (n = 5-7 mice). (D) Specific maximal force of TA muscles expressed as the ratio between regenerating and control legs. Unlike WT, *Bin1*x11-/- muscle force does not increase with time. (E) *Bin1* iso8 mRNA expression in WT mice in non-injured muscle and during regeneration. (F) Fusion index 14 days after notexin injection WT and *Bin1*x11-/- TA muscles. Number of nuclei per fiber was counted on transversal sections. n = 3 mice per group. All graphs depict mean ± s.e.m. (*p < 0.05,**p<0.01,***p<0.001) Student t-test.

To validate the implication of BIN1 muscle-specific isoforms in muscle regeneration, the expression of BIN1 isoforms encompassing exon 11 was investigated by RT-qPCR and found increased at different time points during regeneration (Figure 8E). To investigate potential cell fusion defects that may explain the deficit in early muscle growth during regeneration in the *Bin1*x11-/- mice, nuclei counting was performed in regenerating myofibers and revealed a decreased mean number of nuclei per fiber in *Bin1*x11-/- mice (Figure 8F), correlating with the observed reduction in fiber cross-sectional area and the specific muscle force. In conclusion, BIN1 muscle isoforms appear important for muscle regeneration but not for muscle development.

## Discussion

In *Bin1*x20-/- mice, the SH3 domain present in all isoforms was disrupted, similarly as in some CNM patients with truncating mutations. This led to a strong decrease in BIN1 level and to perinatal death primarily involving muscle defects. Conversely, *Bin1*x11-/- mice, where the muscle-specific isoforms were converted into ubiquitous isoforms by deletion of the in-frame exon 11 coding for the PI domain, had a normal development and muscle function. However, following muscle damage promoted by notexin injection, these mice displayed a delayed muscle regeneration with a defect in myoblast fusion. This study shows that BIN1 is a crucial player in muscle development and that perinatal muscle maturation and function are affected in its absence. In addition, absence of its skeletal muscle-specific isoforms in mice does not alter muscle development but leads to impaired muscle regeneration in adult.

### BIN1 ubiquitous isoforms and skeletal muscle development

Our data confirmed that *Bin1* has the highest expression level in skeletal muscle (Butler et al. 1997; Wechsler-Reya et al. 1997). *Bin1*x20-/- mice die within the first hours after birth. A previously published total *Bin1* KO mice also died at birth, possibly from a suspected cardiomyopathy, with no reported defects of skeletal muscle structure (Muller et al. 2003). However, mice with cardiac-specific BIN1 loss survive to adulthood (Hong et al. 2014; Laury-Kleintop et al. 2015). In our *Bin1*x20-/- model, thorough analysis of heart structure and function revealed no difference with WT. Together with the observation that skeletal muscle specific deletion of exon 20 displays the same lethality as the constitutive deletion, we conclude that postnatal lethality is due to skeletal muscle impairment.

The first *in vivo* study investigating the role of amphiphysins in muscle development was done in drosophila. An amphiphysin null mutant was viable but flightless, and had major T-tubules malformations (Razzaq et al. 2001). There are two amphiphysins in mammals (AMPH1 and BIN1). We show here that mammalian BIN1 has kept a similar function as its drosophila ortholog. In *Bin1*x20-/- mice, the muscle developmental defects are correlated with triads disorganization as well as a major DHPR (T-tubule marker) and RYR1 (SR marker) collapse. These structural defects originate from a lack of T-tubule formation as there was no T-tubules connected to the plasma membrane in *Bin1*x20-/- primary myotubes and as localization of dysferlin and caveolin 3, which are main regulators of T-tubule formation, was impaired in newborn *Bin1*x20-/- muscle. Moreover, we specifically localized BIN1 on triads, in agreement with previous data (Butler et al. 1997). The intracellular tubule network observed by BIN1 immunofluorescence in primary myotubes and isolated myofibers is connected to the membrane, therefore strongly supporting a role for BIN1 in T-tubules formation, as suggested in C2C12 myotubes (Lee et al. 2002). The junctional reticulum defects may be a consequence of the absence of T-tubules, as anchoring of the T-tubule with the SR is essential for maturation of the whole triad and subsequently the muscle fiber (Flucher et al. 1993). In conclusion, BIN1 is not necessary for early myogenesis but is essential for muscle development and perinatal survival through its role on skeletal muscle T-tubule formation. Importantly, we showed BIN1 is also essential for the general organization of myofibers, including mitochondria and nucleus positioning.

### BIN1 muscle-specific isoforms and muscle regeneration

To address if BIN1 has also a role in skeletal muscle maintenance, muscle-specific ablation of *Bin1* was performed in adult at 7 weeks of age using tamoxifen activation of the Cre recombinase under the control of the human skeletal actin promoter. Our data showed that even 25 weeks after the induction of the deletion and although BIN1 protein level showed a strong reduction, the analyzed muscle displayed no difference in histology, ultrastructure or motor function including calcium homeostasis. These data differ from shRNA knockdown of *Bin1* in adult muscle fibers where around 30% of muscle fibers showed swollen T-tubules and greatly reduced Ca^2+^ sparks frequency 14 days after the shRNA electroporation (Tjondrokoesoemo et al. 2011). It is possible that electroporation induced muscle regeneration, which we show to be dependent on BIN1.

The muscle-specific exon 11 is upregulated during muscle differentiation both in vitro and in vivo, and is present in most muscle isoforms in mature murine and human muscle (Butler et al. 1997; Wechsler-Reya et al. 1997; Nicot et al. 2007; Fugier et al. 2011; Cowling et al. 2017). Our *Bin1*x11-/- model did not have significant defects in T-tubule structure at each of the time points tested (2 weeks, 12 weeks and 12 months of age). Our findings are in apparent contradiction with the in vitro data showing that the PI domain is necessary for membrane tubulation in cultured cells (Lee et al. 2002; Nicot et al. 2007; Bohm et al. 2013). Our data indicate that the PI domain is neither required for T-tubule formation, nor muscle development. Similarly, alterations of muscle development in the bin1 zebrafish morphants were rescued with human BIN1 independently of the presence of the PI domain (Smith et al. 2014). Potentially, unidentified regulators of membrane tubulation present in vivo but absent in cultured cells may explain this discrepancy. Among candidates are BIN1 interactors or modifiers of membrane physicochemical properties. Indeed T-tubule defects have been reported in a canine CNM (Walmsley et al. 2017), a model characterized by an altered lipid composition and fluidity of muscle membrane(Blondelle et al. 2015).

Here we reveal a role for the muscle-specific isoforms of BIN1 in muscle regeneration. We used notexin-induced muscle regeneration in *Bin1*x11-/- mice and observed a decrease in fiber crosssectional area at 14 and 28 days after the muscle damage associated with a reduced fusion index and eventually leading to a decrease in specific force recovery. A deficit in myofibers growth during early regeneration was previously linked to a defect in myoblast fusion (Millay et al. 2014; Baghdadi and Tajbakhsh 2018). Additionally, BIN1 was abundant at the myotubes fusion sites (Klinge et al. 2007) and C2C12 cells expressing antisense *Bin1* showed an impaired myoblast fusion (Wechsler-Reya et al. 1998). We cannot exclude a membrane repair component as shRNA-mediated downregulation of *Bin1* compromised myofiber membrane integrity (Tjondrokoesoemo et al. 2011) and as BIN1 was located close to the repair cap (Demonbreun et al. 2016). However, BIN1 was never directly implicated in muscle membrane repair and CK levels were comparable to WT after treadmill exercise in *Bin1*x11-/- mice (757 ± 252 U/l vs. 773 ± 288 U/l). Overall, BIN1 is necessary for muscle development and function, while the PI domain requirement is restricted to muscle regeneration.

### Physiopathology of centronuclear myopathies

In human, *BIN1* mutations lead to autosomal recessive CNM with neonatal onset (Nicot et al. 2007) or autosomal dominant CNM with adult onset (Bohm et al. 2014). Exon 11 skipping causes childhood onset progressive CNM (Bohm et al. 2013), and the mis-splicing of exons 7 and 11 was identified in myotonic dystrophy (Fugier et al. 2011).

*Bin1*x20-/- displayed a strong decrease of BIN1 when its SH3 domain is deleted. Recessive CNM patients have either truncation of the SH3 domain or missense changes in the BAR domain (Nicot et al. 2007; Bohm et al. 2010; Claeys et al. 2010). They exhibit hypotonia at birth and delayed motor development, and several patients died soon after birth. General histology showed highly variable fiber cross-sectional area with a high number of centralized nuclei often clustered, and NADH-TR staining showed oxidative staining aggregations (Romero 2010; Toussaint et al. 2011). The majority of triads had swollen T-tubules or SR, DHPR and RYR1 accumulated in the center of the fiber, disclosing a strong collapse of the T-tubule/SR network. The *Bin1*x20-/- mouse reproduces these histological hallmarks and the muscle impairment, and therefore represents a faithful model to study the mechanisms of recessive CNM. Using this model, we found alteration of the T-tubule formation that could be at the basis of the CNM pathology. BIN1 could potentially regulate DNM2, another protein mutated in CNM, as DNM2 was proposed to modulate T-tubule formation or maintenance in muscle development in drosophila (Chin et al. 2015). Indeed CNM mutations truncating the BIN1 SH3 domain strongly impaired binding to DNM2 (Nicot et al. 2007), and DNM2 downregulation rescued the perinatal death of *Bin1*x20-/- mice (Cowling et al. 2017). Noteworthy the *Bin1*x20+/- heterozygous mice did not display phenotypes, unlike CNM patients with dominant mutations, suggesting dominant CNM is not due to BIN1 haploinsufficiency but rather to a gain-of-function mechanism.

Under unstressed conditions, the *Bin1*x11-/- mice did not develop a CNM phenotype except a slight but significant increase in nuclei mislocalization. Recessive CNM patients were previously identified with a splice site mutation in exon 11, removing the exon 11 whilst not changing the BIN1 level (Bohm et al. 2013). Affected patients had normal speech and motor development. Muscle weakness was noticed at 3.5 years of age, after which the disease became highly progressive and two out of three patients died at the age of 5 and 7 while the third was wheelchair bound. They had a classical CNM histology. *Bin1*x11-/- mice displayed a muscle regeneration defect that could be an important trigger for the patient phenotypes since they were not affected at birth but rather affected by a progressive muscle weakness.

Exon 11 is also mis-spliced in DM, together with abnormal inclusion of brain-specific exon 7 (Fugier et al. 2011). DM is a pleiotropic disease including gradually worsening muscle loss and weakness and a CNM-like histology. Unlike in the *Bin1*x11-/- mice, specific exon 11 skipping using a U7 small nuclear RNA method in adult mice showed a significant reduction of muscle force together with extensive T-tubules disorganization (Fugier et al. 2011). One cannot exclude some U7 off target effects, or conversely a compensatory mechanism during fetal development in the *Bin1*x11-/- mice. Our data suggest that DM muscle phenotypes might be mainly due to exon 7 inclusion rather than exon 11 skipping. Of note, exon 7 inclusion in BIN1 promoted the binding to DNM2 (Ellis et al. 2012), again suggesting that the interplay between BIN1 and DNM2 is important for normal muscle function and in the setup of the CNM and DM myopathies.

In conclusion, BIN1 ubiquitous function is necessary for muscle development and function while BIN1 muscle-specific PI+ isoforms are implicated in muscle regeneration. This supports that BIN1 centronuclear myopathy with congenital onset are due to developmental defects while later onset may be due to regeneration defects.

## Materials and Methods

### Materials

Primary antibodies used were: mouse anti-DHPRα1 (Cav 1.1) subunit (MA3–920; Affinity Bioreagents or ab58552; Abcam), α-actinin (EA-53; Sigma-Aldrich), BIN1 C99D clone (B9428; Sigma-Aldrich), RYR1 (clone 34C; Sigma-Aldrich) DNA polymerase (sc-373884, Santa Cruz) and glyceraldehyde-3-phosphate dehydrogenase (GAPDH, MAB374; Chemicon) monoclonal antibodies; and rabbit anti-RYR1 (a kind gift from Isabelle Marty, Grenoble Institut des Neurosciences, France). Rabbit anti-DNM2 antibodies (R2680 and R2865)(Cowling et al. 2014), anti-MTM1 (R2827)(Hnia et al. 2011) and anti-BIN1 (R2444 against the SH3 domain and R2405 against the PI domain)(Nicot et al. 2007) were made onsite at the antibody facility of IGBMC. Alexa-conjugated secondary antibodies were purchased from Invitrogen. Secondary antibodies against mouse and rabbit IgG, conjugated with horseradish peroxidase (HRP) were purchased from Jackson ImmunoResearch Laboratories. The following products were purchased: Hoechst nuclear stain (B2883, Sigma-Aldrich), ECL chemiluminescent reaction kit (Pierce), LipofectamineTM (Life Technologies), Tamoxifen (Sigma-Aldrich), Notexin (Latoxan).

### Generation of *Bin1* exon 11 and *Bin1* exon 20 deleted mice

The targeting vector was created with LoxP sites flanking exon 11 of *Bin1* or exon 20, then linearized and electroporated into embryonic stem (ES) cells. Recombinant ES cells were injected into C57BL/6 blastocysts that were implanted in pseudo-pregnant females and germline transmission validated. Recombination was triggered using transgenic mice expressing the Cre recombinase under the control of CMV or ACTA1 (human skeletal actin, HSA) promoters. Mice bred and analyzed were B6J strain. In case of time-inducible recombination the *HSACreERt2* transgenic line was used (Miniou et al. 1999). To trigger the recombination, 7-week-old mice were injected daily for 5 consecutive days with 100 μl of tamoxifen solution diluted in 90% vegetable oil and 10% ethanol at 1 mg/100 μl.

### Animal general phenotyping

Animal experimentation was approved by the institutional ethical committee Com’Eth IGBMC-ICS (2012-128). Animals were housed in a temperature-controlled room (19–22 °C) with a 12:12-h light/dark cycle. Mice were weighed weekly until one year of age. Mice were humanely killed when required by CO2 inhalation followed by cervical dislocation, according to national and European legislations on animal experimentation. Mice aged 10-15 weeks were phenotyped under the EUMODIC phenotyping program (www.eumodic.eu) with results made publicly available (www.europhenome.org). Male and female mutant mice were compared to WT littermates. Blood chemistry, echocardiography, Dexascan presented here for male mice (n= minimum 10 per group) were performed as part of pipelines 1 and 2 of the EUMODIC phenotyping program at the Institut Clinique de la Souris (ICS, Illkirch, France; www.ics-mci.fr). Blood glucose concentration was determined with a glucometer and test strips (One touch ultra) both purchased from Lifescan.

### Motor function

String test: Mice are suspended on a wire by their forelimbs, and allowed 20 sec to climb their hindlimbs onto the wire. Three trials per mouse were performed, with 5 min rest between trials. A fall was considered equal to 20 sec (n ≥ 5 mice per group). Grip strength: Performed by placing the 2 front paws or all 4 paws on the grid of a dynamometer (Bioseb, Chaville, France) and pulling mice by the tail in the opposite direction. The maximal strength exerted by the mouse before losing grip was recorded. Three trials per mouse were performed, with 30 sec rest between trials (2 paws test, n ≥ 5 mice per group; 4 paws test, n = 5-7 mice per group). Hanging test: mice were suspended from a cage lid for a maximum of 60 sec. The time the mouse fell off the cage was recorded for each trial. Three trials per mouse were performed. Rotarod test: coordination and whole body muscle strength and fatigability were tested using an accelerated rotating rod test (Panlab, Barcelona, Spain). Mice were placed on the rod which accelerated from 4 to 40 rpm during 5 min. Three trials per day, with 5 min rest between trials, were performed for day 1 (training day) and then for 4 days which were recorded. Animals were scored for their latency to fall (in sec). The mean of the three trials was calculated for each experiment listed above (n = 5-7 mice per group).

### Echocardiography

Pregnant mice were studied on day 17.5-18.5 of gestation (E17.5-18,5) (where 18.5 days is full term), and a total of 19 WT and KO embryos were observed. A vevo 2100 (VisualSonics, Incorporated-Toronto, Canada) system with a 40-MHz transducer (lateral and axial resolutions of 68 and 38 μm, respectively) was used for ultrasound interrogation of the embryos. Pregnant female mice were anesthetized with isoflurane (1.5% isoflurane in medical air containing 21% oxygen) and laid supine in a petri dish filled with physiological solution with the right forearm and left hindlimb implanted with electrocardiogram (ECG) electrodes subcutaneously for heart rate monitoring (450–550 beats/min). Body temperature and physiological solution temperature were both monitored via a thermometer (rectal thermometer for body temperature) and maintained at 36–38 °C using a lamp (for the mouse) and a heating pad (for the saline solution). The obtention of a proper imaging plane involved externalization of the uterus into the warm physiological solution bath followed by placement of the transducer directly on the uterine wall where pre warmed ultrasound gel was applied. M-mode imaging produces images that provided the most accurate measurements of ventricular wall thickness and shortening fraction in fetal mouse heart measuring 2–5 mm. In this mode, the improved frame rate of up to 1,000 frames/sec allowed end systole and end diastole to be measured with ease. At the end of the procedure, the pregnant mouse was killed by cervical dislocation and biopsies of the fetuses were taken for genotyping.

### Muscle contractile properties

*In situ* muscle isometric contraction was measured in response to nerve and muscle stimulation as described previously with a force transducer (Vignaud et al. 2005; Vignaud et al. 2010). Results from nerve stimulation are shown (n = 5-11 mice per group). After contractile measurements, the animals were killed by cervical dislocation. TA muscles were then dissected and weighed to determine specific maximal force.

### *In vivo* muscle regeneration test

12-week-old male *Bin1*x11-/- and WT mice were used. A cycle of degeneration and regeneration was induced and studied in *tibialis anterior* by intramuscular injection of 20 μl notexin diluted in PBS to 10 g/ml in the right leg. The contralateral TA muscle was used as control. At various time-points after the injections, mice were killed by cervical dislocation. The TA muscles of both hindlimbs were immediately removed, snap frozen in isopentane cooled in liquid nitrogen, mounted in OCT, and 8 μm transversal sections were cut and mounted onto SuperFrost+ slides.

### *In Situ* Hybridization

*In situ* hybridizations were performed on whole-mount wild type embryos at E14.5 and E18.5 day (Diez-Roux et al. 2011). Probe used for BIN1 was made by the Eurexpress program (www.eurexpress.org/ee). BIN1 probe sequence:

> TTTTTTTTTTTTTTTTTGGCTTTGGCAGGTTTTTCTTTTTTGTTTGTTTCGCTGCATTTTGAACACTAGGGCTTATT
>
> TTCAAACAGCACACGGTTGGTCTGCAGAGCGGAACCAGGCTGGGCCAGTGTGCAGGCCCTGCCCAGGGCAG
>
> CTGCCAGAAGAGGACCCAAGCCCTGCTCGGTGGCGCAGCCAAGCGTCAGGCAAGTGTGGCGGTGGCTCTGC
>
> ACCCCCGGCCCGCCCTGAACATGCGGGCATCGGGAACTCAACTAGGGGGGGACACAGCAGCTTCAGGAACA
>
> CTGGAGAAGTCCACTGAACGGGGTCGGACGGCTCTTTGGAAAACCACCTCATCTTTGGGGGTATACCTCTTGG
>
> GTTAGGTTTCCGCCCCCCAGTTTCCCGACACTTTTCAAGAATTTCACAACAAAACAGAAACAGAAAAGAGAG

### Primary cell culture

Primary myoblasts from newborn mice were prepared using a protocol adapted from De Palma (De Palma et al. 2010). After hind limb muscles isolation, muscles were minced and digested for 1.5 h in PBS containing 0.5 mg/ml collagenase (Sigma) and 3.5 mg/ml dispase (Gibco). Cell suspension was filtered through a 40 μm cell strainer and pre-plated for 3 h in DMED-10%FBS (Gibco) to discard the majority of fibroblasts and contaminating cells. Non adherent-myogenic cells were collected and plated in IMDM (Gibco)-20% FBS-1% Chick Embryo Extract (MP Biomedical) onto 1:100 Matrigel Reduced Factor (BD Bioscience) in IMDM coated fluorodishes. Differentiation was triggered by medium switch in IMDM + 2% horse serum and 24 h later a thick layer of matrigel (1:3 in IMDM) was added. Myotubes were treated with 80 μg/ml of agrin and the medium was changed every 2 days. For staining membranes accessible to the culture media, myotubes were incubated on ice for 10 min with FM4-64 (5μg/ml; ThermoFisher Scientific) in Ca^2^+ free HBSS before live imaging using a SP8 inverted confocal microscope (Leica Microsystems, Mannheim, Germany).

### Histology on newborns

Day-18.5 fetuses were skinned, fixed in Bouin’s fluid for at least a week and decalcified in rapid decalcifier DC3 (Labonord) 24 h with 2 changes, washed 24 h with 3 changes in 96 % ethanol, dehydrated in absolute ethanol 1 day with 3 changes, cleared in Histolemon 1 day with 3 changes, then embedded in paraplast (4 days with 4 changes). Serial 7 μm thick sections were deparaffinized with Histosol (Shandon) then stained with either Groat’s hematoxylin followed by Mallory’s trichrome or Hematoxylin and Eosin according to standard protocols.

### Histological and immunofluorescence analysis of skeletal muscle

Muscles and other tissues were frozen in nitrogen-cooled isopentane and liquid nitrogen for histological and immunoblot assays, respectively. Longitudinal and transverse 8 μm cryosections of mouse skeletal muscles were prepared, fixed and stained with antibodies to DHPRα 1 (1:100), RYR1 (1:200), α-actinin (1:1,000), DNM2-R2680 (1:200), MTM1-R2827 (1:200), pan-isoform BIN1-C99D (1:50) and BIN1-R2444 (1:100) antibodies. Nuclei were detected by costaining with Hoechst (Sigma-Aldrich) for 10 min. Samples were viewed using a laser scanning confocal microscope (TCS SP5; Leica Microsystems, Mannheim, Germany). Air-dried transverse sections were fixed and stained with hematoxylin and eosin (HE), succinate dehydrogenase (SDH), NADH-TR or Sirius red/fast green staining and image acquisition performed with a slide scanner NanoZoomer 2 HT equipped with the fluorescence module L11600-21 (Hamamatsu Photonics, Japan) or a DMRXA2 microscope (Leica Microsystems Gmbh). Cross-sectional area (CSA) was analyzed in HE sections from TA skeletal muscle using FIJI image analysis software. CSA (μm^2^) was calculated (> 500 fibers per mouse) from 4-7 mice per group. The percentage of TA muscle fibers with centralized or internalized nuclei was counted in > 500 fibers from 4-6 mice using the cell counter plugin in ImageJ image analysis software.

### Immunofluorescence on isolated muscle fibers

Dissected muscles were fixed in 4% PFA for 30 min at RT, and then incubated in PBS supplemented with 0.1 M glucose for 30 min at RT and afterwards in PBS supplemented with 30% sucrose at 4 °C overnight. Muscles were then frozen at -80 °C. Thawn entire muscles were dissected in PBS to isolate fibers. Fibers were permeabilized in PBS supplemented with 50 mM NH4Cl and 0.5% Triton X100 for 30 min at RT and then saturated in PBS supplemented with 50 mM NH4Cl, 0.5% donkey serum and 0.1% Triton X100 for 1 h at RT. Immunofluorescence was performed at 4 °C overnight using MTM1-R2827 (1:200), BIN1 (1:50, C99D), DHPR (1:150, Abcam) and RYR1 (1:150, Sigma-Aldrich) antibodies. Fibers were then incubated with donkey anti-mouse or donkey anti-rabbit secondary antibodies (Alexa Fluor 488/594, Life technologies) diluted 1:250.

### Light microscopy

All microscopy was performed at the IGBMC Imaging Centre. All samples for microscopy were mounted in Fluorsave reagent (Merck) and viewed at room temperature. Light microscopy was performed using a fluorescence microscope (DM4000; Leica microsystems) fitted with a color CCD camera (Coolsnap cf colour, Photometrics). Confocal microscopy was performed using a confocal laser scanning microscope (TCS SP2 or SP5; Leica Microsystems, Mannheim, Germany). ImageJ and FIJI analysis software were used for image analysis (Schneider et al. 2012).

### Transmission electron microscopy

Mice were anesthetized by intraperitoneal injection of 10 μl/g of ketamine (20 mg/mL; Virbac) and xylazine (0.4%, Rompun; Bayer) and euthanized. Muscle biopsy specimens from hind limbs were fixed with 2.5% glutaraldehyde in 0.1 M cacodylate buffer (pH 7.2) and processed as described (Amoasii et al. 2012). For T-tubule analysis, potassium ferrocyanide staining was performed as described (Al-Qusairi et al. 2009).

### Immunogold staining

Deeply anaesthetized mice were transcardially perfused with 4% formaldehyde, 0.1 M phosphate buffer. Dissected *tibialis anterior* were further post-fixed in 4% formaldehyde and cut in 100 μm thick longitudinal sections with a vibratome. A standard free-floating immunocytochemical procedure was followed, using 0.1 M saline phosphate buffer as diluent and rinsing liquids. After preincubation in 5% normal goat serum and 5% BSA, sections were incubated overnight at 4 °C in 1/500 anti BIN1-2405 antibody. A further 4 h incubation with ultra-small gold conjugate of goat antimouse IgG (1/20; Aurion, Netherlands) was followed by extensive washings, 10 min post-fixation in 2% glutaraldehyde and 0.70 nm gold beads were then silver enhanced (HQ silver; Nanoprobes, Stony Brook, NY). After 15 min post-fixation in 1% OsO4, sections were dehydrated in graded acetone and finally embedded in Epon resin. Ultrathin sections were examined with a Philips CM120 electron microscope, operated at 80kV and imaged with a SIS Morada digital camera.

### RT-PCR and quantitative RT-PCR

Total RNA was purified from hind limb, TA and quadriceps muscles using TRIzol reagent (Invitrogen) according to manufacturer’s instructions. cDNA was synthetized from 1–2 μg of total RNA using SuperScript II reverse transcriptase (Invitrogen) and random hexamers. Quantitative PCR amplification of cDNA was performed on a Light-Cycler 480 instrument (Roche). Gene expression is considered as dysregulated when fold change is higher than 1.3 and p value is ≤ 0.05. Following primers were used (from 5’-3’):

> Bin1 exon7 F: ACTATGAGTCTCTTCAAACCGCC
>
> Bin1 exon7 R: TCCACGTTCATCTCCTCGAACACC
>
> Bin1 exon9 F: TCAACACGTTCCAGAGCATC
>
> Bin1 exon19 R: GTGTAATCATGCTGGGCTTG
>
> *Bin1* exon18-19 F : CATGTTCAAGGTTCAAG
>
> *Bin1* exon20 R: TGATTCCAGTCGCTCTCCTT
>
> *Casq* 2 F: GCCCAACGTCATCCCTAACA
>
> *Casq* 2 R: GGGTCACTCTTCTCCGCAAA
>
> *Tnnt* 2 F: GCCATCGACCACCTGAATGA
>
> *Tnnt* 2 R: GCTGCTTGAACTTTTCCTGC
>
> *Jp* 2 F: ACGGAGGAACCTATCAAGGC
>
> *Jp* 2 R: CTTGAGAAAGCTCAAGCGGC
>
> *eMhc* F: AGATGGAAGTGTTTGGCATA
>
> *eMhc* R: GGCATACACGTCCTCTGGCT
>
> *Gapdh* F: TTGTGATGGGTGTGAACCAC
>
> *Gapdh* R: TTCAGCTCTGGGATGACCTT

### Western blotting

Muscle was homogenized using an Ultra-Turrax (IKA-WERKE) homogenizer in 50 mM Tris, 10% glycerol, 50 mM KCl, 0.1% SDS, 2% Triton X 100 and a set of protease inhibitors: 1 mM EDTA, 10 mM NaF, 1 mM Na3VO4, 1 mM PMSF (phenylmethylsulphonyl fluoride), 1 μM pepstatin, 10 μM leupeptin. The homogenate was kept at 4 °C for 2 h, and clarified by centrifugation at 13000 rpm for 10 min. Protein concentration of the supernatant fraction was quantified with the Biorad Protein Asssay (Biorad laboratorie GmbH) and lysates analyzed by SDS-PAGE and western blotting on nitrocellulose membrane. Primary antibodies used were DNM2-R2680 (1:500), DNM2-R2865 (1:500), BIN1-R2405 (1:5000), BIN1-R2444 (1:500), GAPDH (1:10,000) and DNA polymerase (1:1000); secondary antibodies were anti-rabbit HRP or anti-mouse HRP (1:10,000). Western blot films were scanned and band intensities were determined using ImageJ software. Densitometry values were standardized to corresponding total GAPDH values and expressed as a fold difference relative to the listed control (n = 3-5 mice per group).

### Electrophysiology and fluorescence measurements in isolated muscle fibers

Single fibers were isolated from the *flexor digitorum brevis* (FDB) and interosseus muscles following previously described procedures (Jacquemond 1997). Comparisons were always made between groups of fibers issued from the same muscle type. Experiments were carried out on muscle fibres isolated from the muscles of both hindlimbs from 3 WT and 3 *Bin1*x11-/- mice.

The silicone voltage-clamp technique (Jacquemond 1997; Lefebvre et al. 2014) was used with an RK-400 patch-clamp amplifier (Bio-Logic, Claix, France) in combination with an analog-digital converter (Digidata 1440A, Axon Instruments, Foster City, CA) controlled by pClamp 9 software (Axon Instruments). Voltage-clamp was performed with a micropipette filled with a solution mimicking the ionic composition of the cytosol and also containing either a high concentration of EGTA and the fluorescent Ca^2+^ -sensitive probe rhod-2, or the contraction blocking agent N-benzyl-p-toluene sulphonamide (BTS) and the fluorescent Ca^2+^ -sensitive probe fluo-4 FF (see Solutions). Intracellular equilibration of the solution was allowed for 30 min. For calcium current analysis, linear leak component was removed as previously described (Kutchukian et al. 2017). The voltage dependence of the peak current was fitted with the following equation: *I*(*V*) = *G_max_*(*V-V_rev_*)/[1+exp(*V_0.5_-V*)/*k*] with *I(V)* the peak current density at the command voltage *V*, *G_max_* the maximum conductance, *V_rev_* the apparent reversal potential, *V_0.5_* the half-activation potential and *k* the steepness factor.

Confocal imaging was conducted with a Zeiss LSM 5 Exciter microscope equipped with a 63× oil immersion objective (numerical aperture 1.4). For detection of rhod-2 and fluo-4 FF fluorescence, excitation was from the 543 nm line of a HeNe laser and from the 488 nm line of an Argon laser, respectively, and fluorescence was collected above 560 nm and above 505 nm, respectively. Rhod-2 and fluo-4 FF Ca^2+^ transients were imaged using the line-scan mode (*x*,*t*) of the system. Rhod-2 and fluo-4 FF fluorescence changes were expressed as F/F_0_ where F_0_ is the baseline fluorescence. Quantification of the spatial heterogeneity of Ca^2+^ release activation and of the absolute Ca^2+^ release flux underlying the rhod-2 Ca^2+^ transients were performed as previously described (Kutchukian et al. 2017). In each fiber, the voltage-dependence of the peak rate of Ca^2+^ release was fitted with a Boltzmann function. For imaging the T-tubule network, FDB muscle fibers were incubated for 30 minutes in the presence of 10 μM di-8-anepps in Tyrode solution. For Ca^2+^ sparks measurements, FDB fibres were incubated for 30 minutes in the presence of Tyrode solution containing 10 μM fluo-4 AM. For both di-8-anepps and fluo-4 imaging, fluorescence was collected above 505 nm with 488 nm excitation. The T-tubule density from the di-8-anepps fluorescence was estimated as described previously (Kutchukian et al. 2017).

Tyrode solution contained (in mM): 140 NaCl, 5 KCl, 2.5 CaCl_2_, 2 MgCl_2_, 10 HEPES. The extracellular solution used for voltage-clamp contained (in mM) 140 TEA-methanesulfonate, 2.5 CaCl_2_, 2 MgCl_2_, 1 4-aminopyridine, 10 HEPES and 0.002 tetrodotoxin. The pipette solution contained (in mM) 120 K-glutamate, 5 Na_2_-ATP, 5 Na_2_-phosphocreatine, 5.5 MgCl_2_, 15 EGTA, 6 CaCl_2_, 0.1 rhod-2, 5 glucose, 5 HEPES. All solutions were adjusted to pH 7.2.

### Statistical analysis

Since the compared groups came from the same background, statistical analyses were performed using the unpaired student’s T-test unless stated otherwise. p-values of < 0.05 were considered significant.

## Author contributions

I. P., J.Lap. designed the study. I.P., B.S.C., C.K., C.Kr., H.T., J.H., O.W., A.F., A.T., C.G., V.N., C.Ko., J.Lai., R.C., V.J. performed research. L.T. provided scientific advice. I.P., B.S.C., F.S., J.Lai., R.C., V.J., F.P-S., J.Lap. analyzed data. B.S.C., J.Lap. supervised the study. I.P. and J.Lap. wrote the manuscript which was edited by L.T. and F.P.S. and approved by all co-authors.

## Supporting information

## Acknowledgements

We thank Stéphanie Gadin, Aymeline Vandestienne and Alexandre Prola for experimental help, Vincent Gache for advices and IGBMC and ICS institutional platforms of genetic engineering, phenotyping, animal care, histology, photonic and electron imaging for excellent technical assistance. This study was supported by institute funding from INSERM, CNRS, University of Strasbourg and by grants from the Agence Nationale de la Recherche ANR-08-GENOPAT-005, ANR-10-LABX-0030-INRT under the frame program Investissements d’Avenir ANR-10-IDEX-0002-02, Réseau National des Génopoles (RNG), Fondation Maladies Rares, Fondation pour la Recherche Médicale (20071210538), and Association Française contre les Myopathies (13304, 14058, 15352 and Translamuscle).

