## Supplementary material for "Differential physiological role of BIN1 isoforms in skeletal muscle development, function and regeneration"

### Supplementary materials

**Supplementary Figure 1. *BIN1* expression and targeted disruption of mouse *Bin1* exon 20.** (A) *BIN1* RNA levels in adult human tissues based on the GTEx database; TPM = transcripts per million. (B) The genomic region surrounding the targeted exon 20 of *Bin1* in mice; L3: construct inserted into the locus, and L-: deleted allele. (C) PCR from genomic DNA of WT, *Bin1*x20-/- and *Bin1*x20+/- mice. (D-E) Constitutive deletion was achieved with a CMV promoter and induced a strong reduction in total BIN1 protein level in skeletal muscle. (D) Western blot with an antibody against the muscle-specific PI domain. (E) No compensatory upregulation of AMPH1 using anti-amphiphysin 1 antibody. (F) Skeletal muscle deletion was achieved with the ACTA1 (human skeletal actin, HSA) promoter and induced a strong reduction in total BIN1 protein level in skeletal muscle.

**Supplementary Figure 2. Heart characterization in constitutive *Bin1*x20-/- mice.** (A) Total body weight of newborn WT and *Bin1*x20-/- mice was not significantly different (n ≥ 8 mice per group). (B) Ratio of the heart weight on body weight (n ≥ 7 mice per group). (C) Histology of the heart in a newborn WT and two *Bin1*x20-/- mice; 2 serial sections shown. Scale bar 200 µm. (D) Echocardiography done in utero on E18.5 embryos showing no difference in the systolic (DTS) and diastolic (DTD) contraction in left (upper panel) or right (lower panel) ventricle as well as in heart rate (n ≥ 8 mice per group). (E) RT-qPCR from the heart of newborn P0 mice showing no difference in relative expression of calsequestrin 2 (CASQ2), troponin T (TNNT2) and junctophilin 2 (JP2), three markers of cardiac integrity (n ≥ 3 mice per group).

**Supplementary Figure 3. Anomalies of mitochondria positioning in *Bin1*x20-/- muscles.** (A) NADH-TR and SDH staining in newborn WT and KO mice showing collapse of staining towards the center in KO muscle. Scale bar 200 µm. (B) Prohibitin localization on KO and WT 8 µm transversal muscle sections at birth. Scale bar 10 µm.

**Supplementary Figure 4. Normal localization of several sarcomeric markers in newborn constitutive *Bin1*x20-/- mice.** (A) α-actinin and DNM2 stainings on 8 µm transversal muscle sections. Scale bar 10 µm. (B) DNM2 staining on isolated fibers. Scale bar 5 µm. Note that muscle fibers in WT newborn are usually less polygonal than in adult.

**Supplementary Figure 5. Phenotyping of adult *Bin1*x20+/- heterozygous mice at 12 weeks.** (A) Body weight of *Bin1*x20+/- mice (n = 5) and wild type (WT, n = 6) littermates. (B) Muscle mass percent in WT and *Bin1*x20+/- mice. (C) Grip strength and (D) rotarod performance in female and male *Bin1*x20+/- mice. (E) HE staining of TA muscles. Scale bar 50 µm. (F) Potassium-ferrocyanide staining of T-tubules and intracellular ultrastructure of *Bin1*x20+/- and WT muscles. Scale bar 1 µm.

**Supplementary Figure 6. Tamoxifen induced *Bin1* exon 20 deletion from 7 weeks of age.** (A) The genomic region surrounding the targeted exon 20 of *Bin1*; L2: floxed allele and L-: deleted allele. (B) PCR showing exon 20 excision from muscles of constitutive *Bin1*x20-/-, tamoxifen treated control (floxed L2 *Bin1*x20 without Cre recombinase) and tamoxifen treated L2 *Bin1*x20skm(i) (thus *Bin1*x20skm(i)-/-) mice. (C) RT-qPCR from floxed *Bin1*x20 controls and *Bin1*x20skm(i)-/- mice from *tibialis anterior* (TA) and quadriceps (Q) Five weeks after tamoxifen injections. (D) BIN1 protein level 5 weeks after intraperitoneal tamoxifen injection. All graphs depict mean ± s.e.m. (*$p < 0.05$). Student t-test; n ≥ 3 mice per group.

**Supplementary Figure 7. Phenotyping of *Bin1*x20skm(i)-/- mice 25 weeks after tamoxifen injection.** (A) Body weight. Control is tamoxifen treated floxed L2 *Bin1*x20 without Cre recombinase. (B) Ratio of muscle/body weight of different muscles (TA, *tibialis anterior*; Q, quadriceps; EDL, *extensor digitorum longus*; SOL, *soleus*). (C) The grip test was performed using the 2 front paws. (D) String test (time taken to rise hind limbs to wire). (E) Hanging test (time suspended from a cage lid, max time 60 sec). n ≥ 6 mice per group.

**Supplementary Figure 8. Targeted disruption of mouse *Bin1* exon 11 to create *Bin1*x11-/- mice.** (A) The genomic region surrounding the targeted exon 11 of *Bin1*. Deletion of the muscle-specific exon 11 is predicted to lead to an in-frame transcript corresponding to ubiquitous isoforms. (B) PCR genotyping from the WT (+/+), *Bin1*x11-/- and *Bin1*x11+/- mice. (C) RT-PCR from WT and *Bin1*x11-/- TA muscles between exons 9 and 19 showing complete removal of exon 11 in *Bin1*x11-/- and the presence of 2 isoforms differing by inclusion or exclusion of alternative exon 17. *Bin1* iso8 (+/- exon 17) contains the exon 11 and is found only in muscle while *Bin1* iso9 (+/- exon 17) does not encompass exon 11 and is ubiquitous. (D) cDNA sequences from WT and *Bin1*x11-/- TA muscles, showing exon 11 is removed and neighboring exons are present. (E) Western blot using muscle-specific BIN1 antibody (anti-PI domain) showing complete lack of the PI domain encoded by exon 11 in TA. Pan-BIN1 antibody (against SH3 domain) showing the presence of iso9. (F) Western blot from brain and TA muscle probed with anti-AMPH1 antibody.

**Supplementary Figure 9. Body composition of 12-week-old *Bin1*x11-/- mice.** (A) Body weight and (B) body length of *Bin1*x11-/- mice did not differ from WT littermates. (C) DEXA scan analysis did not show any strong difference in the body composition between *Bin1*x11-/- mice and WT. (D) Muscle mass over body weight ratio for *tibialis anterior* (TA), *extensor digitorum longus* (EDL) and *soleus* (SOL). All graphs depict mean ± s.e.m (n ≥ 7 mice per group).

**Supplementary Figure 10. Muscle function analysis in 12-week-old *Bin1*x11-/- mice.** The Grip test was performed using 2 (A) or 4 (B) paws. (C) Further motor evaluation was performed with rotarod and the latency before mice felt was noted. (D) The specific maximal force of the TA muscle represents the absolute maximal force related to muscle weight. All graph depict mean ± s.e.m (n ≥ 6 mice per group).

**Supplementary Figure 11. Histological and ultrastructural analyses of *Bin1*x11-/- muscles at 12 weeks.** (A) TA muscles fiber area; fiber cross-sectional area is grouped into 200 µm² intervals, and represented as the percentage of total fibers (n = 5-7 mice). (B) Frequency of fibers with internal or central nuclei (n = 5 mice). Internal nuclei are defined as not subsarcolemmal nor central. (C) TA muscle ultrastructure analyzed by electron microscopy. Scale bar 1 µm. All graph depict mean ± s.e.m. (*p < 0.05, **p < 0.01). Student t-test.

**Supplementary Figure 12. DHPR calcium current in muscle fibers from WT and *Bin1*x11-/- mice**. Representative $Ca^{2+}$ current traces from a fiber isolated from a WT mouse (A) and from a *Bin1*x11-/- mouse (B) in response to the pulse protocol shown above. (C) mean voltage-dependence of the peak $Ca^{2+}$ current in the two groups of fibers. (D) Mean values for the parameters obtained from fitting each current *vs* voltage dataset with Equation 1 (see methods).

**Supplementary Figure 13. Voltage-activated SR calcium release in muscle fibers from WT and *Bin1*x11-/- mice.** Representative rhod-2 $Ca^{2+}$ transients ($F_0$) and below corresponding $Ca^{2+}$ release flux traces ($d[Ca_{Tot}]/dt$) in a muscle fiber from a WT mouse (A) and from a *Bin1*x11-/- mouse (B) in response to the pulse protocol shown on top. Dotted graphs present the mean voltage-dependence of the peak rate of SR $Ca^{2+}$ release (C) and of the time top peak $Ca^{2+}$ release (D) in the two groups of muscle fibers. (E) Mean values for maximal rate of SR $Ca^{2+}$ release (Max rate), mid-activation voltage ($V_{0.5}$) and steepness factor (k) obtained in the two groups of fibers, by fitting a Boltzmann function to data from each fiber.

**Supplementary Figure 14. Myoplasmic $Ca^{2+}$ removal in muscle fibers from WT and *Bin1*x11-/- mice.** Fluo-4 FF $Ca^{2+}$ transients elicited by short voltage-clamp depolarizing pulses of increasing duration in a muscle fiber from a WT (A) and from a *Bin1*x11-/- mouse (B). (C) Mean values for the rate constant of $Ca^{2+}$ decay in the two groups of fibers, obtained from fitting a single exponential function to the declining phase of the transients after the end of the pulses.

**Supplementary Figure 15. Body composition analysis at 12 months of age in *Bin1*x11-/- mice.** (A) Body weight of *Bin1*x11-/- mice was slightly increased compared to WT. (B) The increase in body weight did not come from an increase in muscle mass. (C) qNMR analysis of *Bin1*x11-/- mice confirmed that lean mass was normal but fat mass was slightly increased and normalized free body fluid (FBF) value was slightly decreased. (D) Normalization of the individual muscles mass compared to the lean tissue. (E) Specific maximal force of the TA muscle represents the absolute maximal force related to muscle weight. (F) TA muscles fiber area; fiber cross-sectional area is grouped into

200 µm² intervals, and represented as the percentage of total fibers (n = 5-7 mice). All graph depict mean ± s.e.m. (*p < 0.05, **p < 0.01) (n ≥ 7 mice per group). Student t-test.

**Supplementary Figure 16. Muscle organization in 12-month-old mice.** (A) Isolated fibers from EDL were stained with BIN1 (pan-isoform antibody), DHPR or RYR1 antibodies. Scale bar 10 µm. (B) Isolated fibers from EDL were stained with titin (green) and α-actinin (red) antibodies. Scale bar 10 µm. (C) Sarcomere organization of TA assessed by electron microscopy. Scale bar 1 µm. Sarcomere and triad organization is similar between *Bin1*x11-/- and WT littermates.

**Supplementary 17. Characterization of *Bin1*x11-/- mice at 2 weeks of age during postnatal hypertrophy.** (A) Body weight. (B) Muscle weight normalized to body weight for quadriceps (Q) and *tibialis anterior* (TA). (C) Histological analysis with hematoxylin and eosin staining. Scale bar 100 µm. (D) TA fiber area grouped into 200 µm² intervals, and represented as the percentage of total fibers. N = 5-6 mice per group. (E) Frequency of fibers with internal or central nuclei. Internal nuclei are defined as not subsarcolemmal nor central. (F) Ultrastructural analysis of TA muscles and T-tubules through potassium ferrocyanide labeling. Scale bar 1 µm.

**Supplementary Figure 18. Muscle regeneration induced by notexin injection in TA muscles of 12-week-old WT and *Bin1*x11-/- mice.** Histology of the muscles stained with hematoxylin and eosin (HE) and observed at 5, 7, 14 and 28 days after the notexin injection. Scale bar 200 µm.

A

| BIN1 | Skel. muscle | Cardiac muscle | Brain | Liver | Arthery | Bladder | Nerve | Kidney | Pancreas | Spleen | Testis |
|---|---|---|---|---|---|---|---|---|---|---|---|
| Median TPM | 935,4 | 7,7 | 144,2 | 17,9 | 28,1 | 124,3 | 143,2 | 62,6 | 11,9 | 82,1 | 22 |

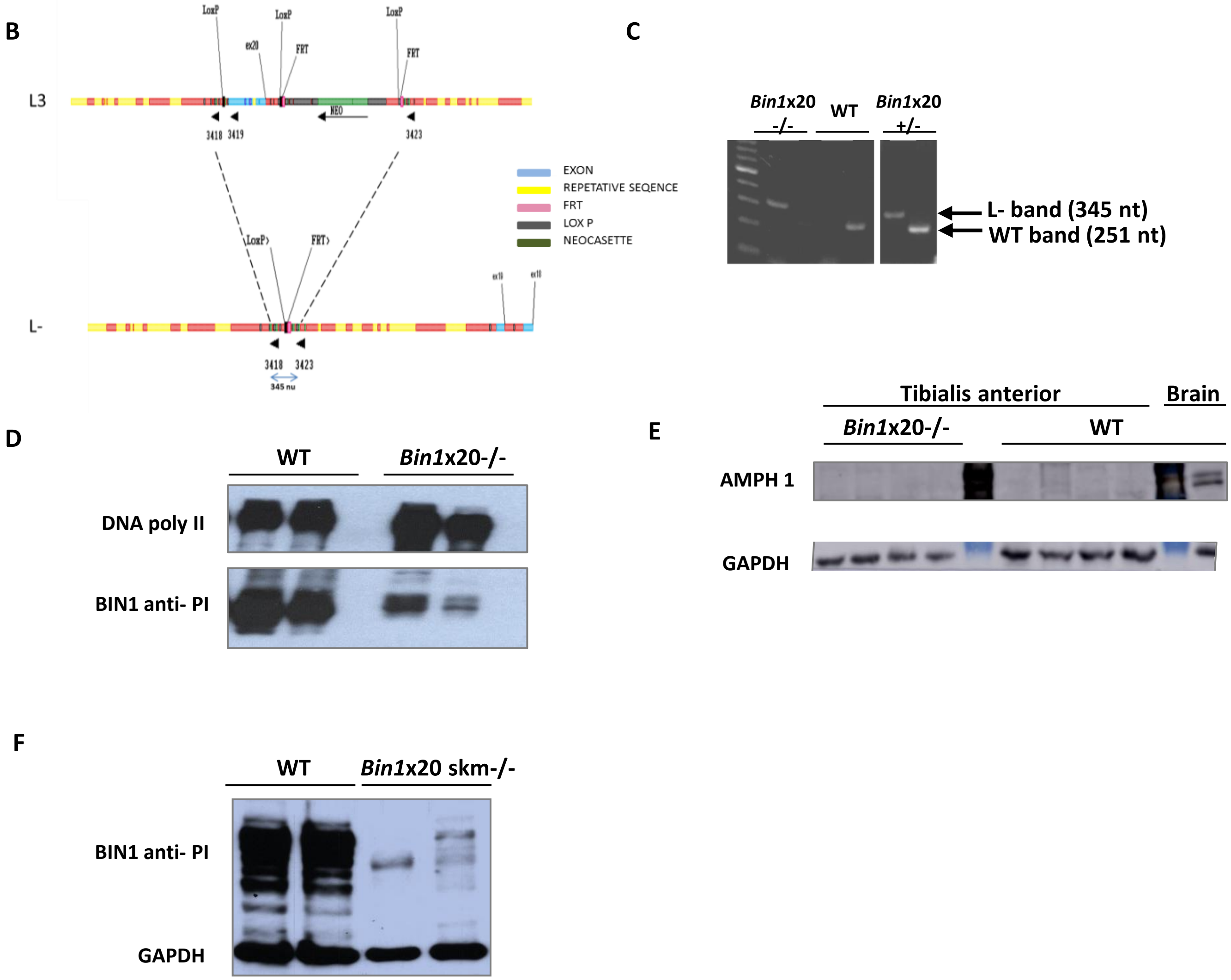


**Supplementary Figure 1.**

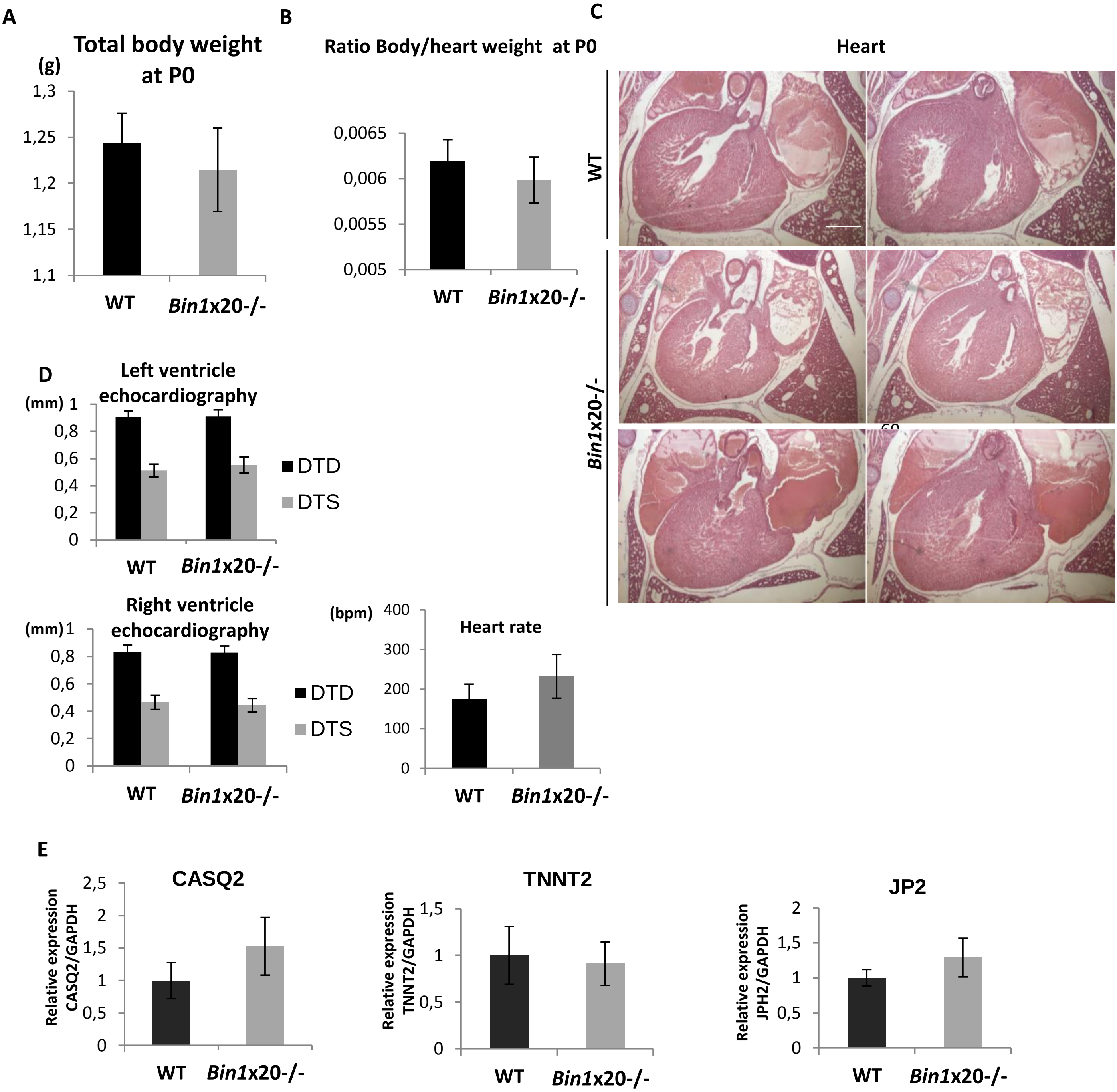


**Supplementary Figure 2.**

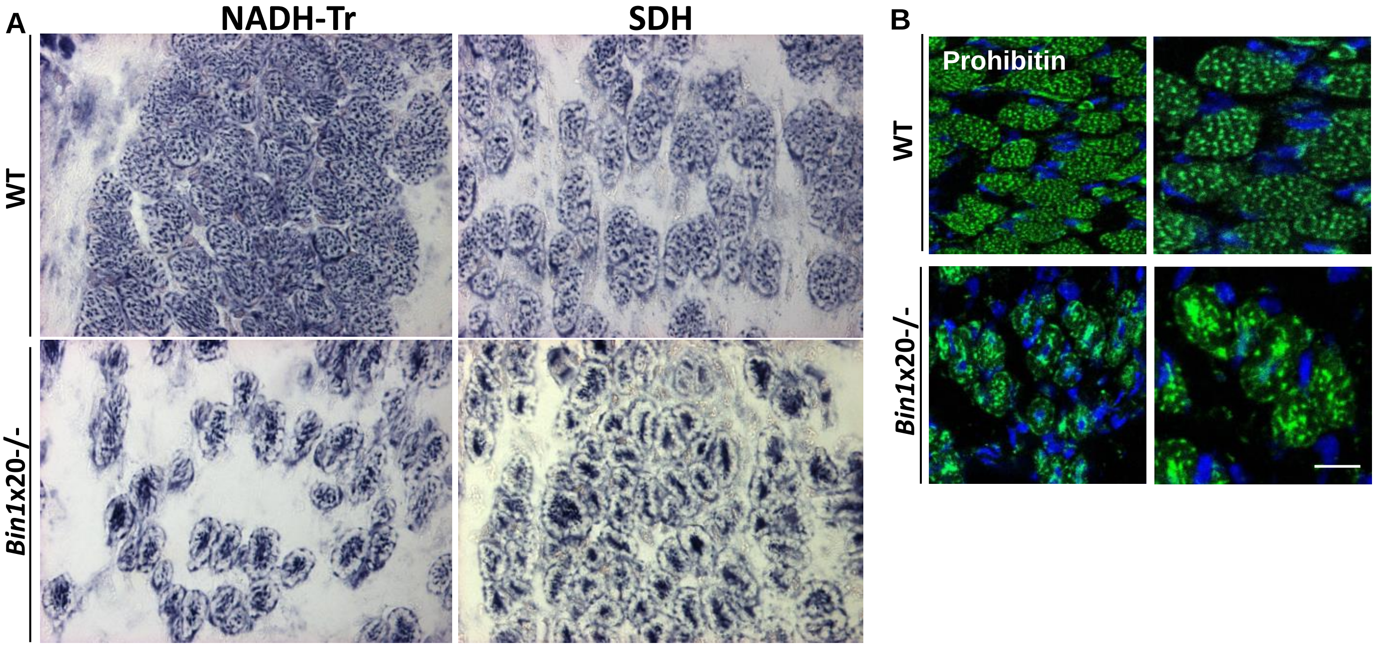


**Supplementary Figure 3.**

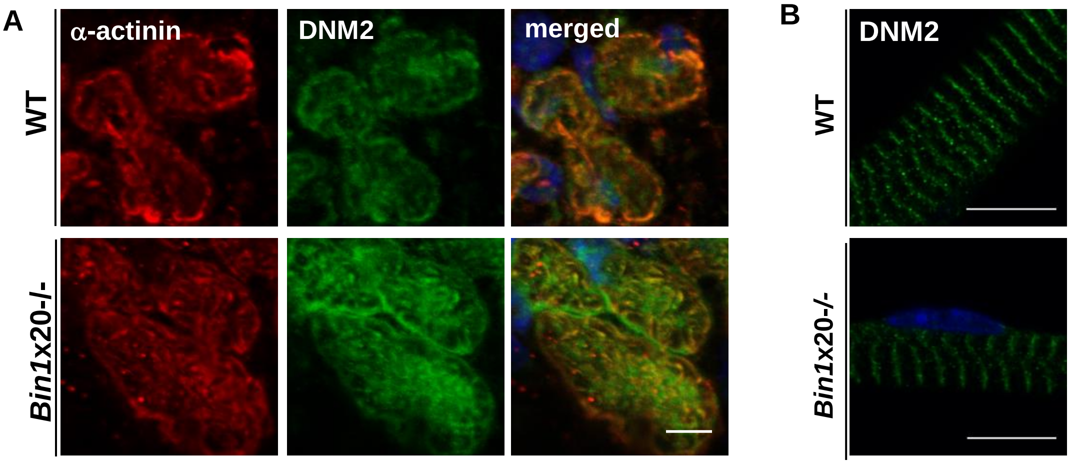


**Supplementary Figure 4.**

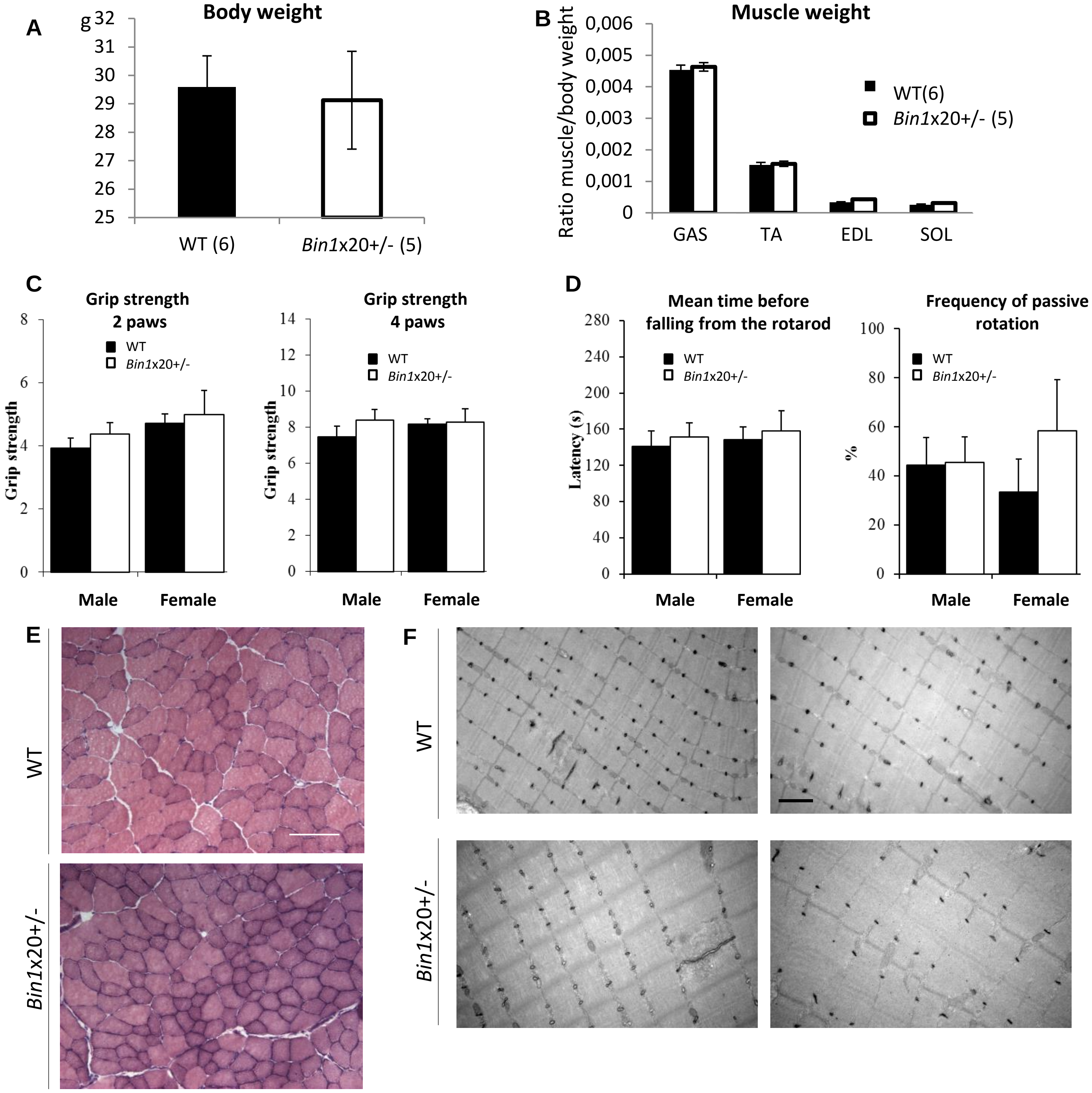


**Supplementary Figure 5.**

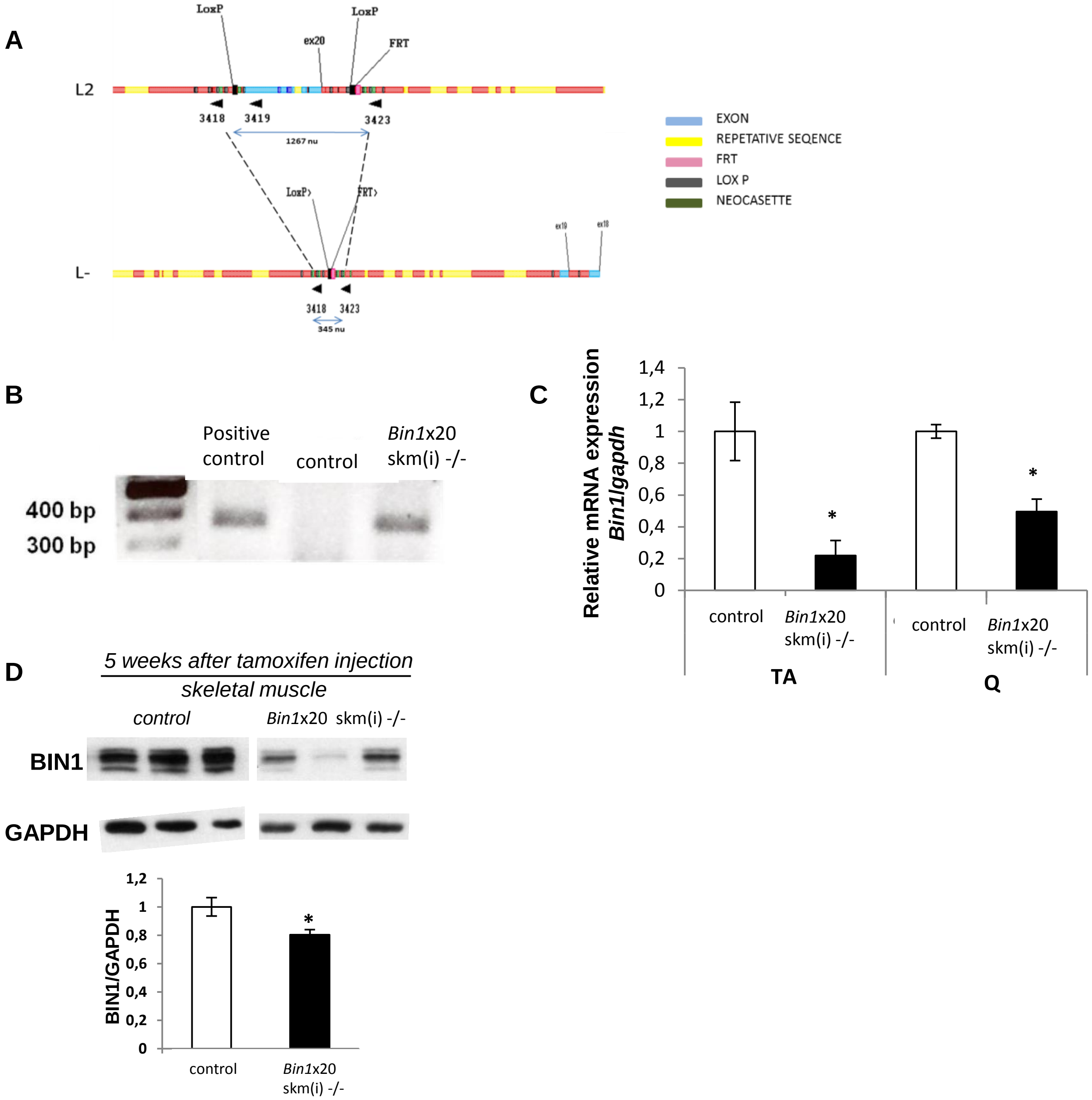


**Supplementary Figure 6.**

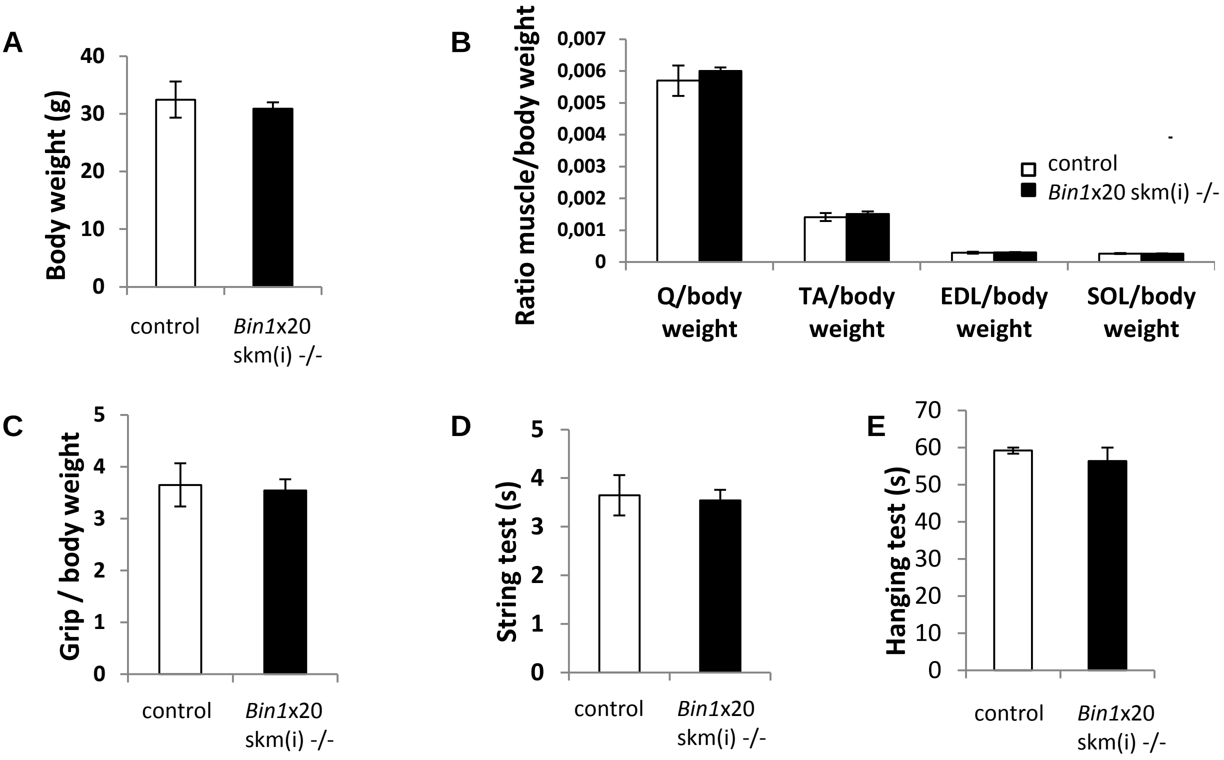


**Supplementary Figure 7.**

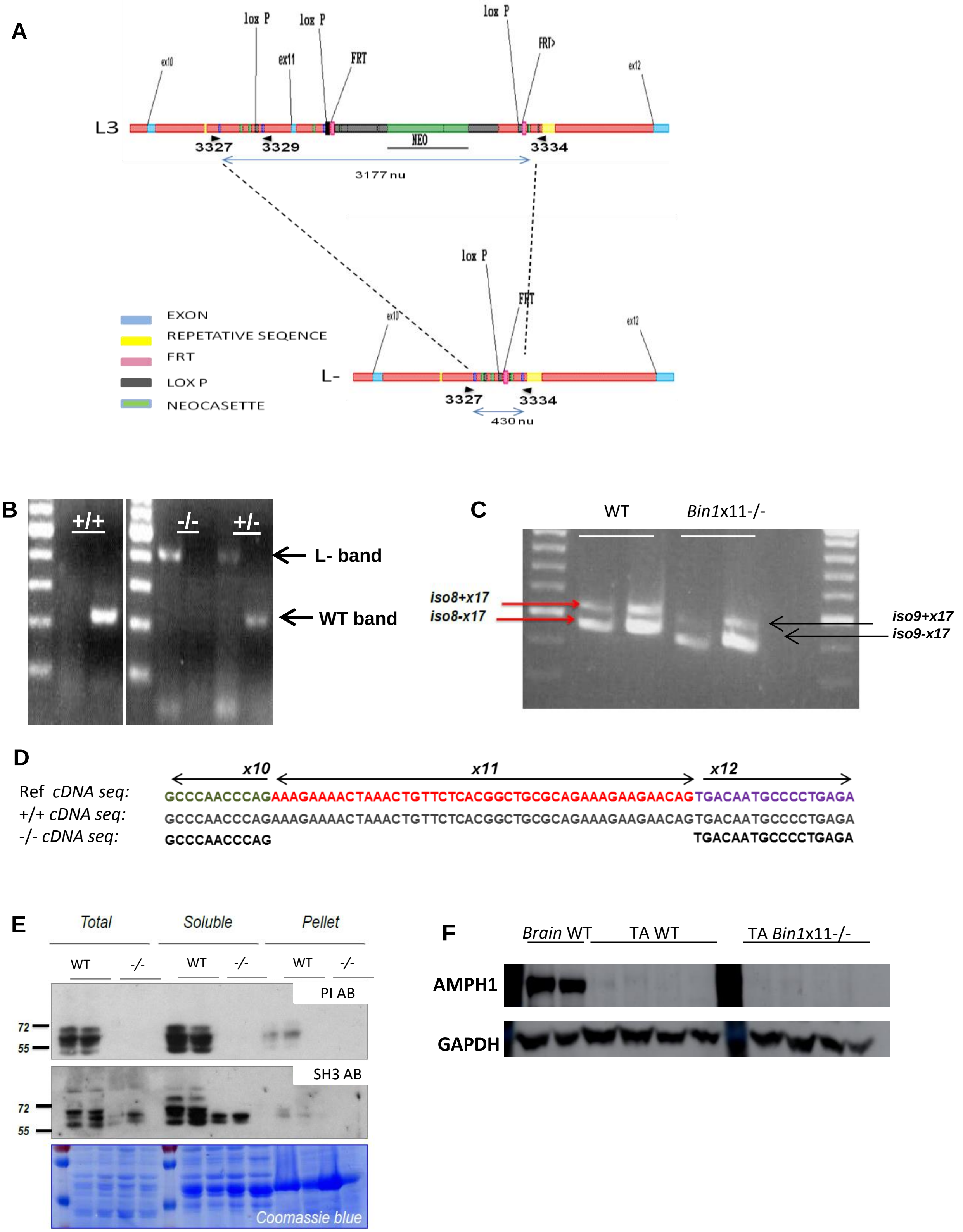


**Supplementary Figure 8.**

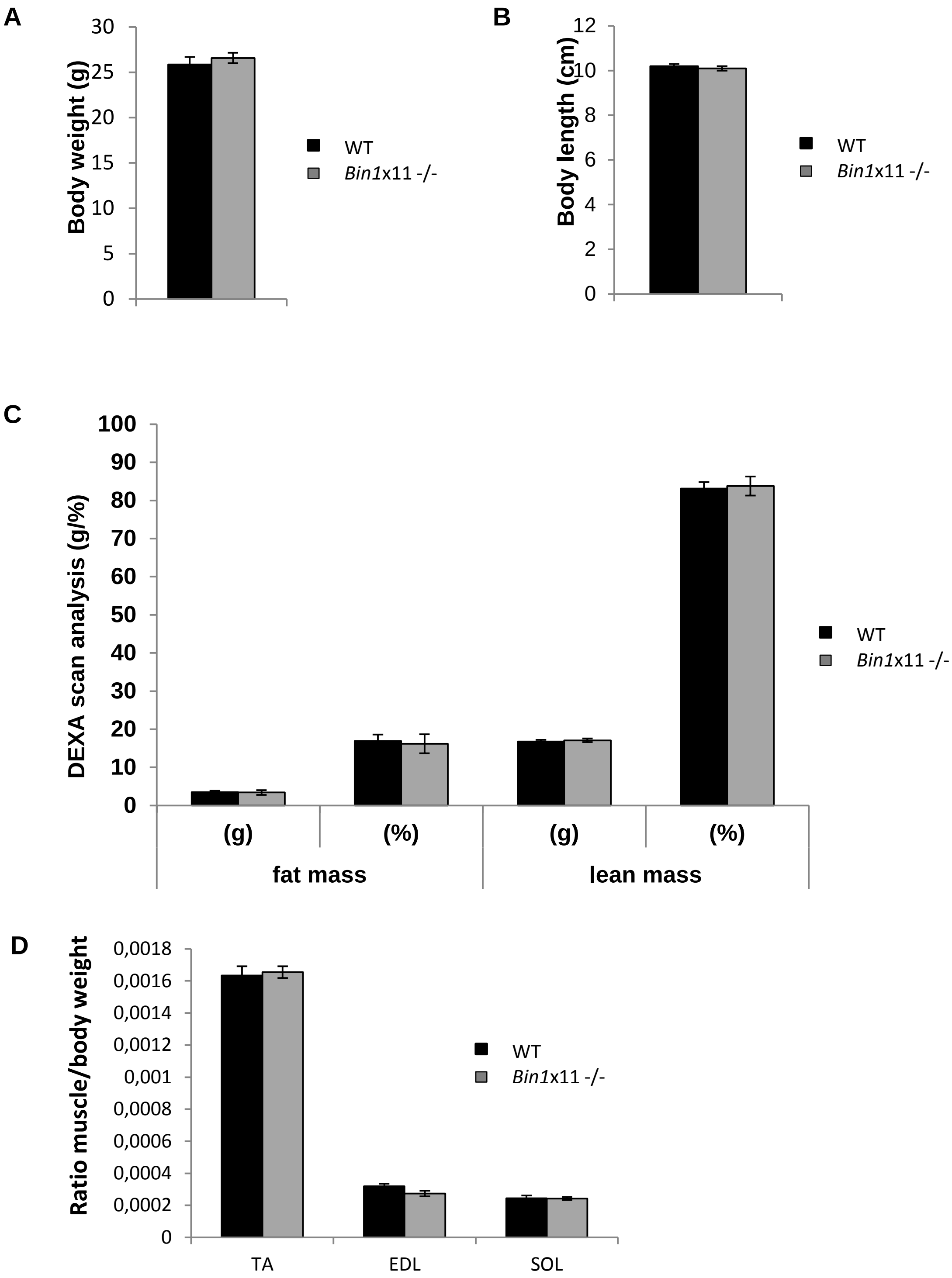


**Supplementary Figure 9.**

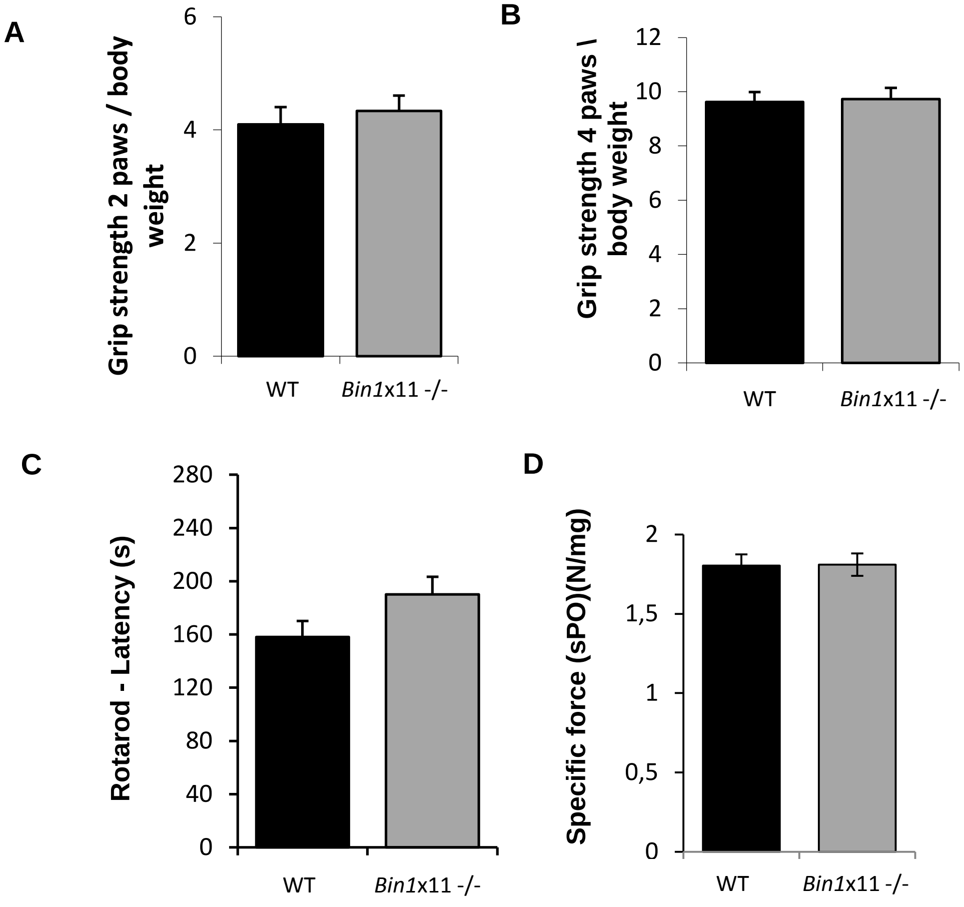


**Supplementary Figure 10.**

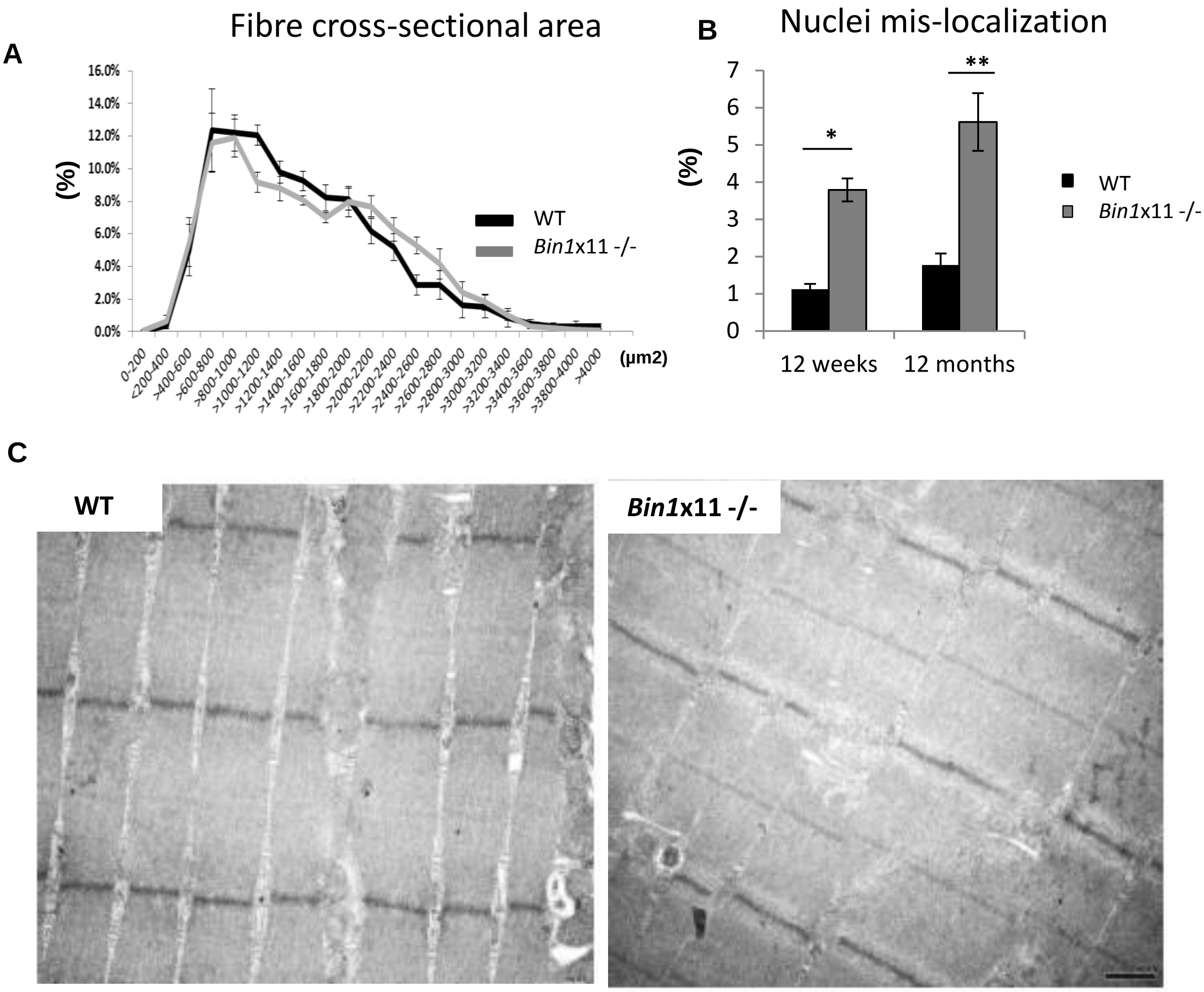


**Supplementary Figure 11.**

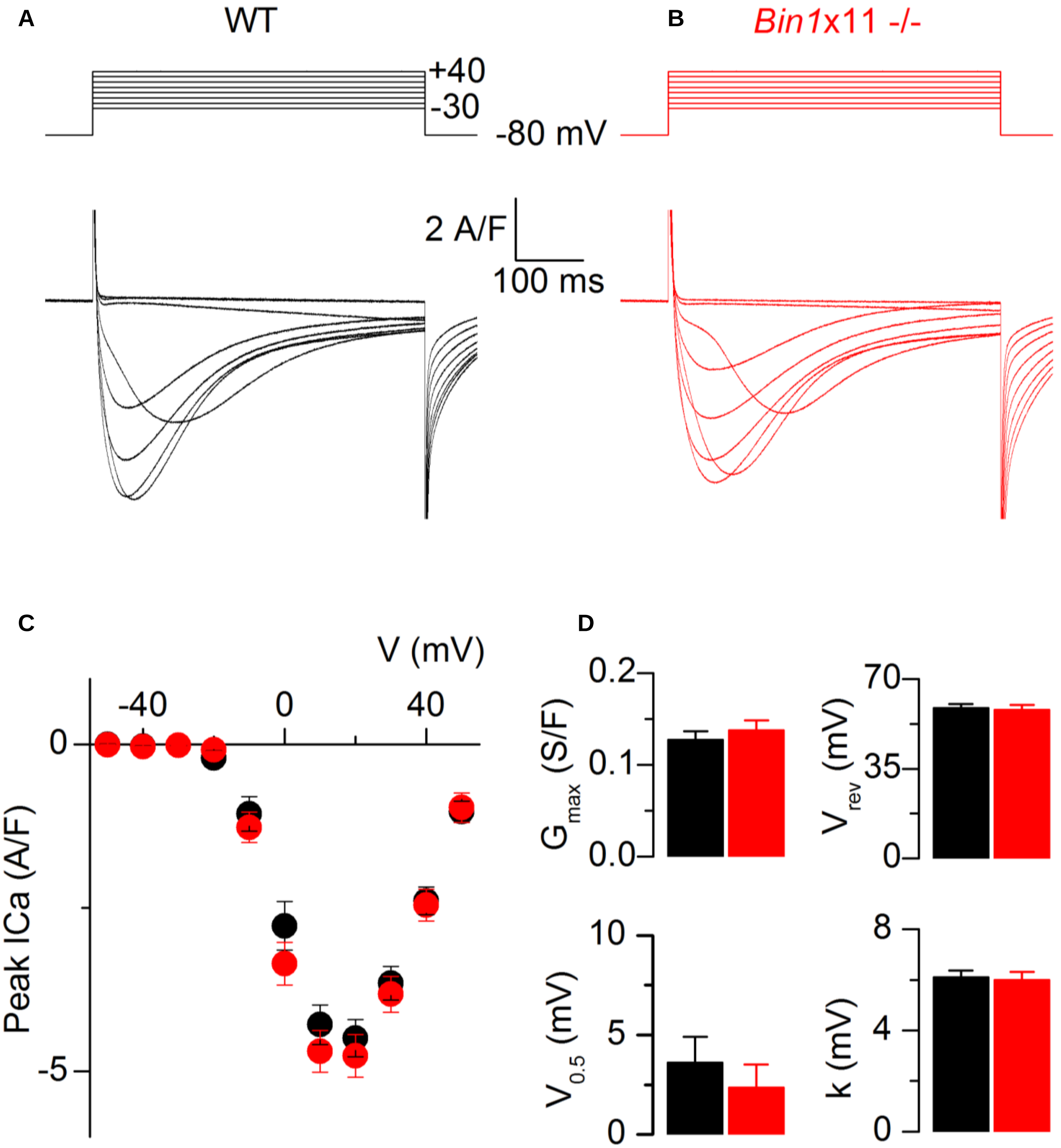


**Supplementary Figure 12.**

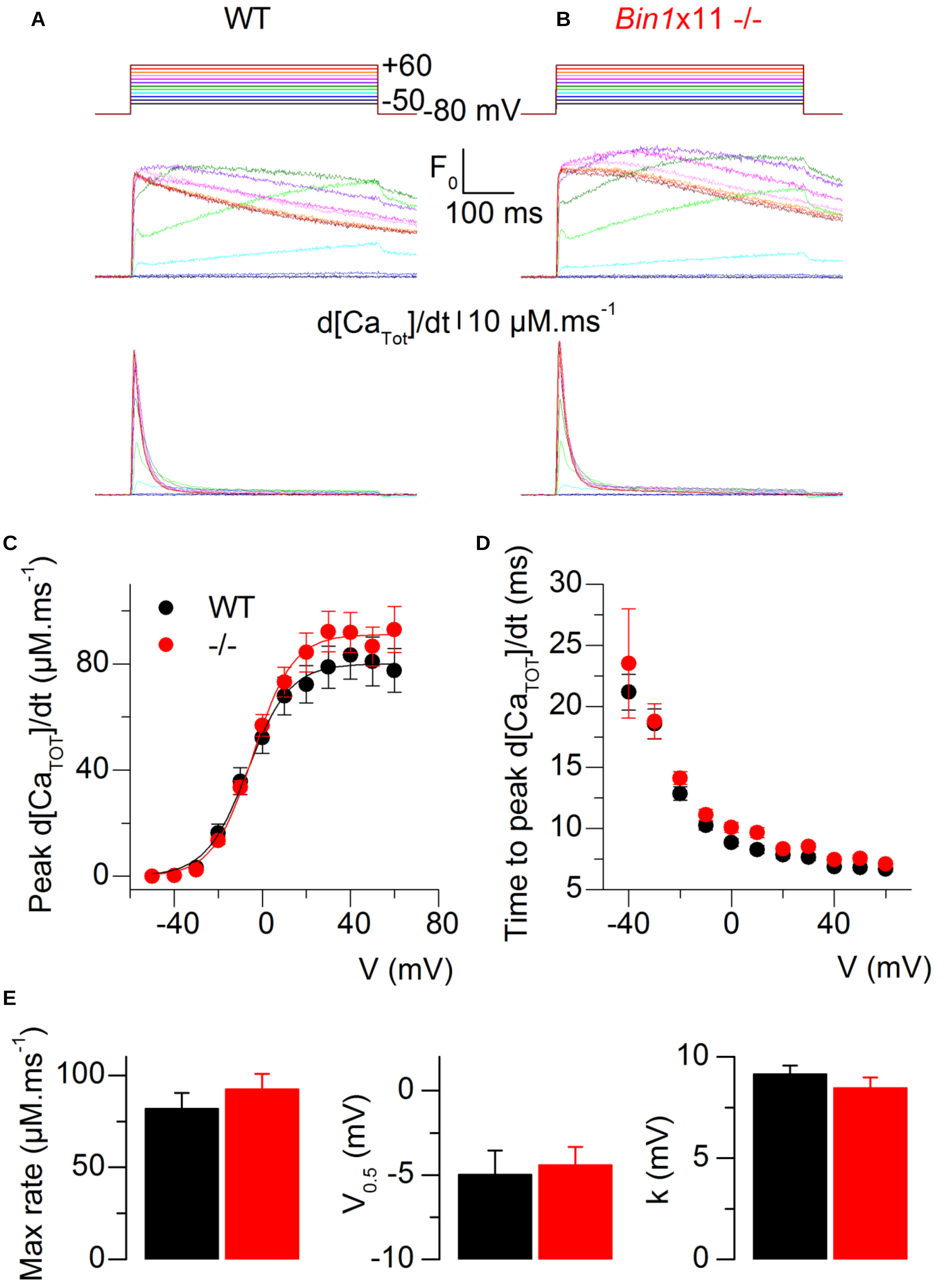


**Supplementary Figure 13.**

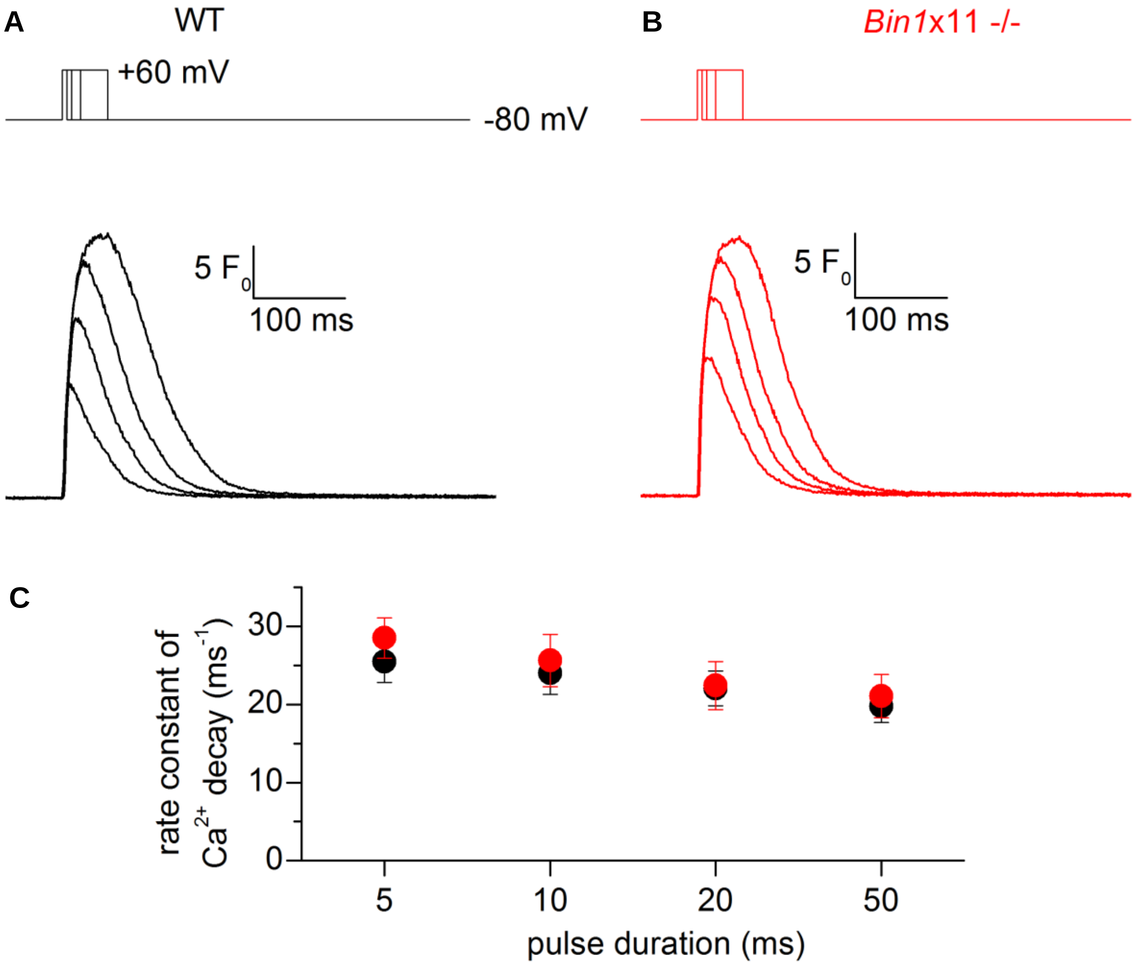


**Supplementary Figure 14.**

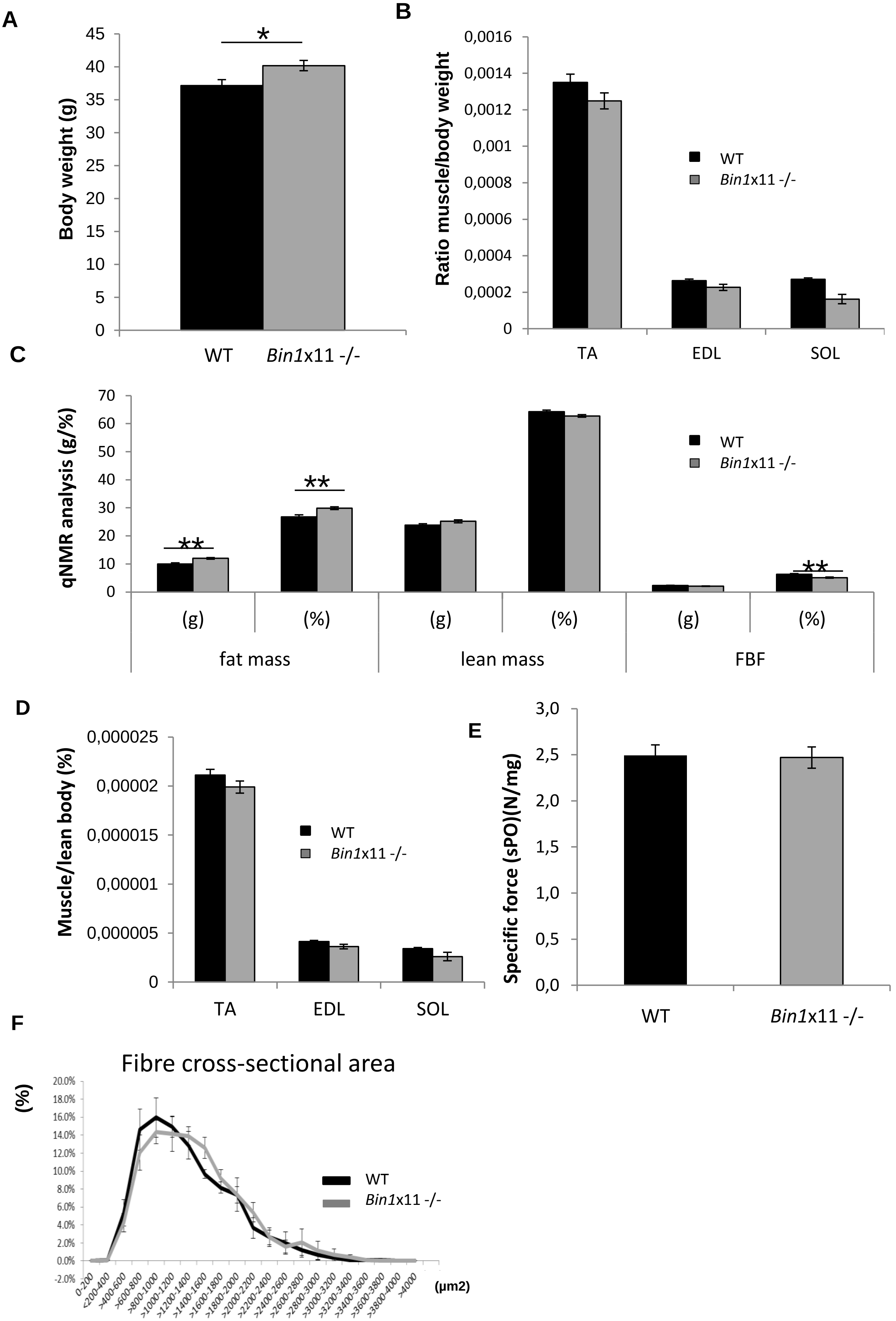


**Supplementary Figure 15.**

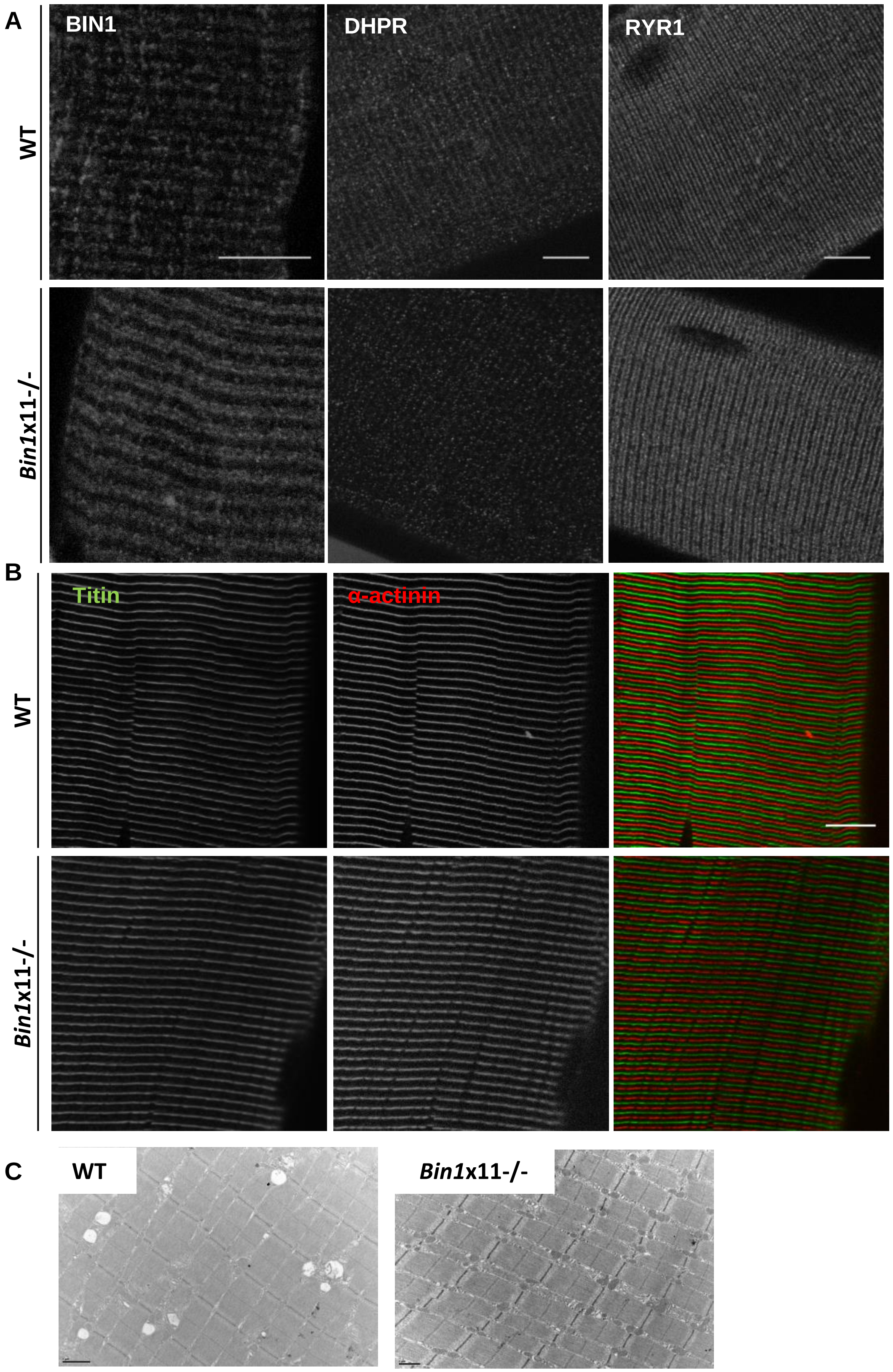


**Supplementary Figure 16.**

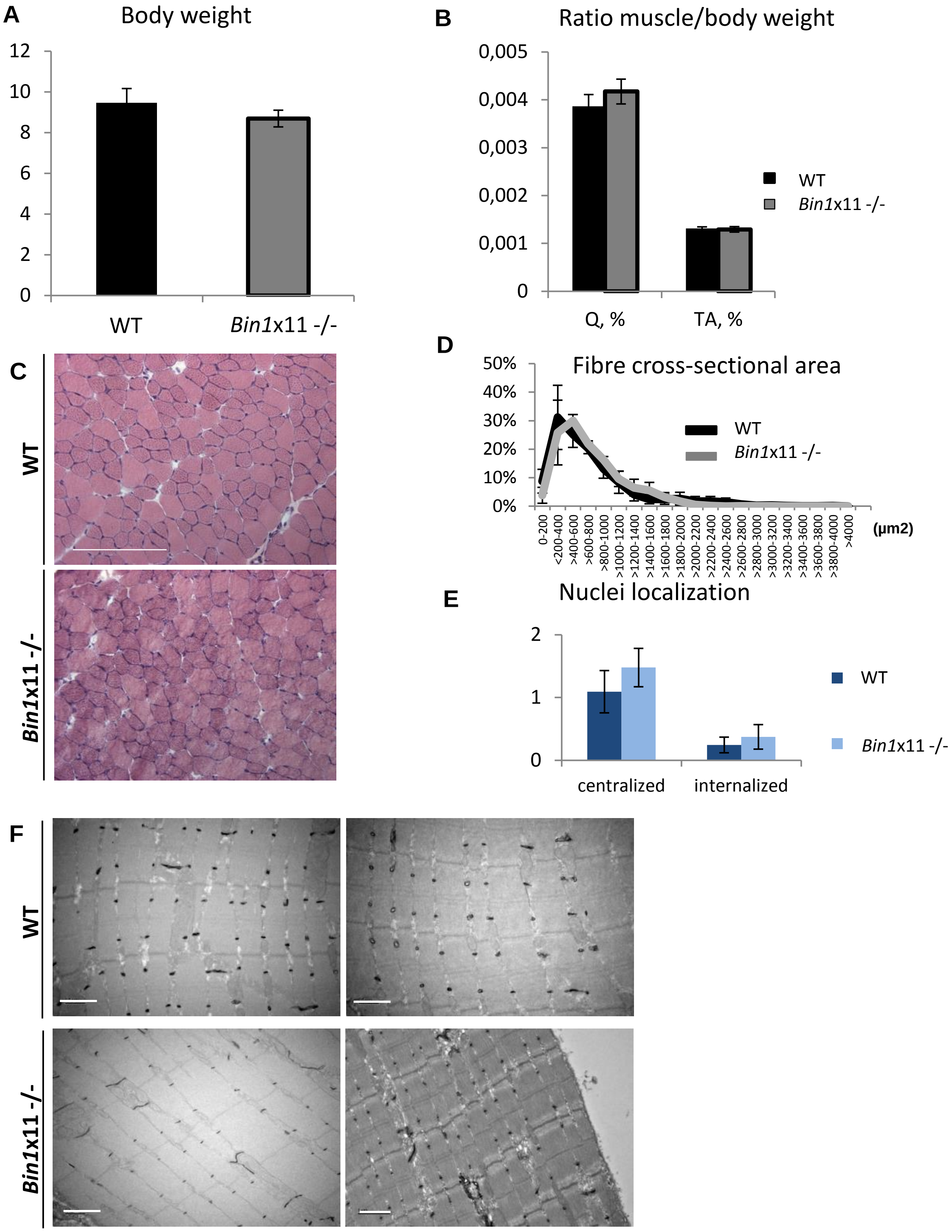


**Supplementary Figure 17.**

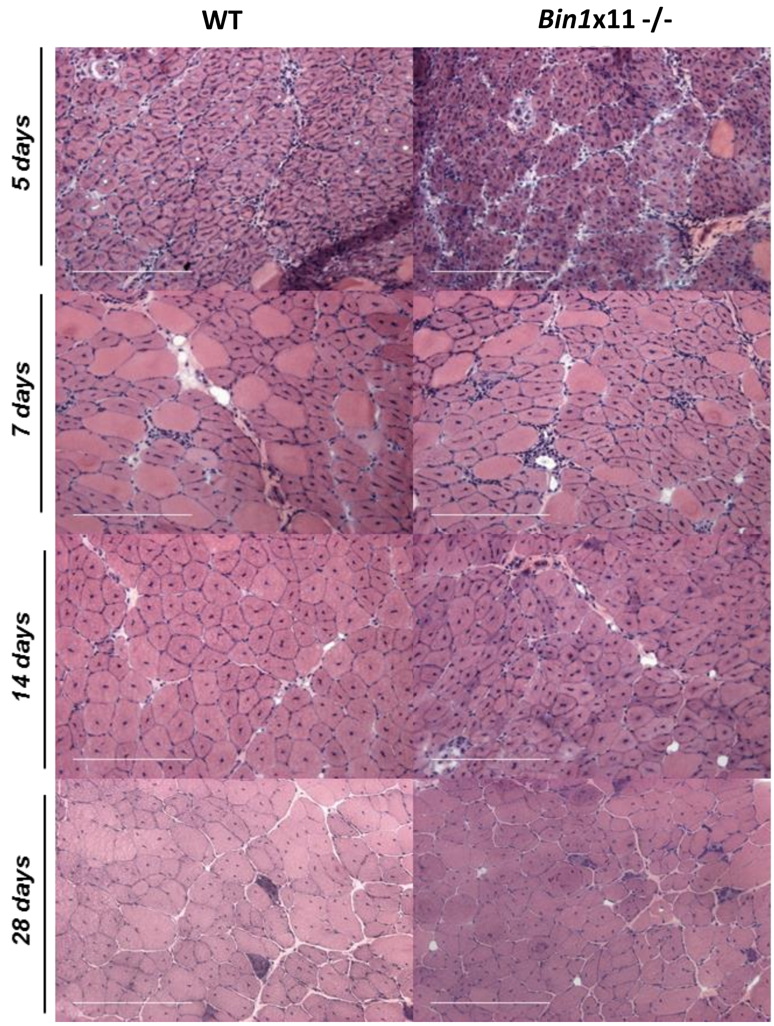


**Supplementary Figure 18.**
